# Probabilistic Retinotopic Parcellation of the Macaque Visual Cortex

**DOI:** 10.64898/2026.07.23.740239

**Authors:** Nora E. Fitzgerald, Qi Zhu, Wim Vanduffel

## Abstract

Probabilistic brain atlases provide anatomically and functionally interpretable normative references that summarize population-level organization while explicitly representing spatial uncertainty. Yet such resources remain scarce due to the resource-intensive experimental efforts required to construct them. Here, we present a probabilistic retinotopic atlas of the macaque visual cortex, derived from contrast-enhanced phase-encoded fMRI data acquired in 13 subjects. The dataset includes a 50% probability parcellation covering 19 visual areas, individual subject labels, and voxel-wise probability maps for each area, all registered to the MEBRAINS macaque template. By combining a consensus parcellation with spatial estimates of confidence for each visual area, this atlas enables more informed anatomical localization and interpretation than deterministic or single-subject-based atlases. As a standardized reference for the macaque visual cortex, it supports experimental design, data interpretation, and multimodal data integration while providing a quantitative framework for investigating the developmental and evolutionary principles that shape the primate visual cortex.

## Introduction

Retinotopic organization, or the orderly mapping of visual space onto subcortical nuclei or the cortical surface, is a defining feature of the primate visual system. Neighboring neurons respond to neighboring locations in the visual field, collectively forming representations that span much or all of the contralateral visual hemifield and, in some regions, extend into the ipsilateral hemifield^1–6^. Consequently, retinotopic organization provides one of the principal criteria for delineating visual cortical areas, an essential task as about half of the primate cortex is devoted to visual processing^7,8^. Yet, despite decades of investigation, the organization of the visual cortex remains incompletely resolved. New retinotopically defined areas continue to be identified^9–11^, established areas continue to reveal greater organizational complexity^12^, and the boundaries of many visual areas remain debated because of methodological limitations, limited sample sizes, and varying mapping and delineation criteria^2,13–18^.

One source of these discrepancies is the conventional strategy used to construct atlases based on retinotopically defined visual areas in non-human primates (NHPs). Typically, data from only a few animals are registered to a common anatomical reference space, often derived from a single subject, and merged into a consensus parcellation^9,13,19,20^. While these atlases have provided invaluable references for systems neuroscience, they necessarily obscure inter-individual and inter-hemispheric variation in the location and extent of visual areas and assign discrete areal boundaries without quantifying confidence or uncertainty^6^. Their interpretation is further complicated by the coexistence of functional and architectonic parcellation schemes. Many widely used macaque brain atlases are based on cytoarchitectonic features, including acetylcholinesterase reactivity and immunoreactivity for parvalbumin, neurofilament protein SMI-32, tyrosine hydroxylase, and other molecular markers^21,22^. The extent to which these complementary frameworks converge remains an open question, highlighting the need for population-level functional references that can be directly compared with architectonic atlases.

Probabilistic atlases address these limitations by estimating, for every location in a common reference space, the probability of belonging to a given area across a population, rather than assigning deterministic boundaries^6,23^. In doing so, they capture both consensus cortical organization and the uncertainty arising from inter-individual variability and imprecise areal borders. Such atlases have transformed human brain mapping and are increasingly being developed for the macaque cortex using anatomical and functional datasets^6,24–31^. Beyond providing richer anatomical references, probabilistic atlases improve the planning and interpretation of electrophysiological recordings, viral injections, optical imaging, functional ultrasound, and other spatially constrained methodologies while enabling quantitative analyses of cortical variability and allowing meta-analyses. However, generating reliable probabilistic atlases requires substantially larger cohorts than conventional atlas studies, making such resources particularly scarce for *in vivo* NHP studies.

Here, we present the first large-scale probabilistic atlas of retinotopically defined visual areas in the macaque cortex, generated from fMRI data acquired in thirteen subjects. Alongside probabilistic areal maps, we provide group-average polar angle and eccentricity maps derived from the same dataset, allowing direct visualization of the underlying retinotopic organization. Registered to the MEBRAINS macaque template^32^, these resources establish a standardized population reference that explicitly quantifies uncertainty in visual area localization and retinotopic organization. By combining probabilistic areal assignments with group-average retinotopic maps, this resource facilitates multimodal studies of the macaque visual system, enables direct comparisons between functional and architectonic parcellations, and provides a quantitative framework for investigating inter-individual variability and the developmental and evolutionary principles that shape primate visual cortex.

## Results

### Acquisition and Average Representations

We acquired phase-encoded retinotopic data from 13 awake rhesus macaque subjects during a series of behaviorally controlled fMRI experiments. To maximize responses across retinotopically organized areas, we used two highly salient stimulus sets consisting of dynamic monkey faces and walking humans (Figure 1A)^29^. Phase-encoded time series were analyzed using Fourier transforms to estimate eccentricity and polar angle responses in each subject^2^. These maps were transformed from their native brain space to the MEBRAINS template via the F99 template. Averaging the resulting phase-maps across subjects and converting them to visual degrees yielded fine-grained average eccentricity and polar angle maps (Figure 1B) spanning the occipital cortex and extending into early parietal cortex up to 12.5° eccentricity. Surface representations revealed a clear and precise retinotopic organization not only throughout early visual cortex, but also in the posterior bank of the superior temporal sulcus (STS) as well as portions of the intraparietal sulcus (IPS).

**Figure 1:**
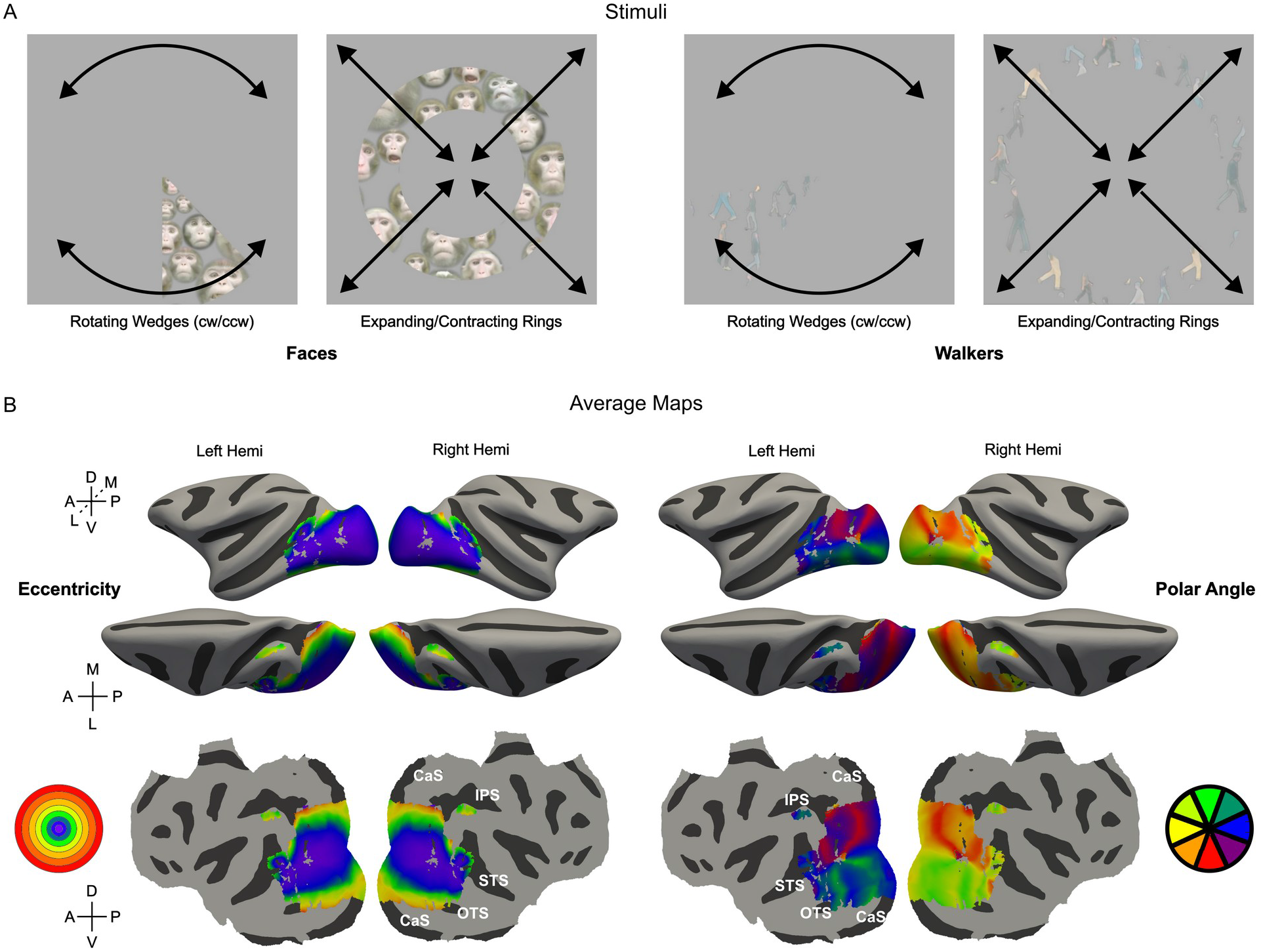
Phase-encoded Retinotopic Stimuli and Average Eccentricity/Polar Angle Maps (A) Two types of stimuli were shown: ‘Faces’ and ‘Walkers’ in both rotating wedge (clockwise and counterclockwise) and expanding/contracting ring formats. Black motion arrows are representative, not part of the displayed stimuli. (B, left) Average eccentricity map (visual angle) across subjects shown on inflated surface and flatmap representations of the MEBRAINS template for both left and right hemispheres. Color coding is shown in circular map. (B, right) Same for polar angle. Sulci names are shown in white

Although these average maps were not used for subsequent analyses, they provide a useful reference for evaluating the probabilistic parcellation and for visualizing the overall organization of retinotopic representations across the visual cortex. Figure 2 shows the average eccentricity and polar angle maps projected onto left and right hemisphere flatmaps of the MEBRAINS template together with iso-eccentricity and iso-polar angle contours (Figure 2B/D) and the probabilistic parcellation (Figure 2C). Iso-eccentricity lines were plotted at 1°, 2°, 4°, and 8°, while iso-polar angle contours divided the visual field into 16 equal angular sections (Figure 2B). As expected, central field representations (< 6° eccentricity) occupied much of the occipital visual cortex.

**Figure 2:**
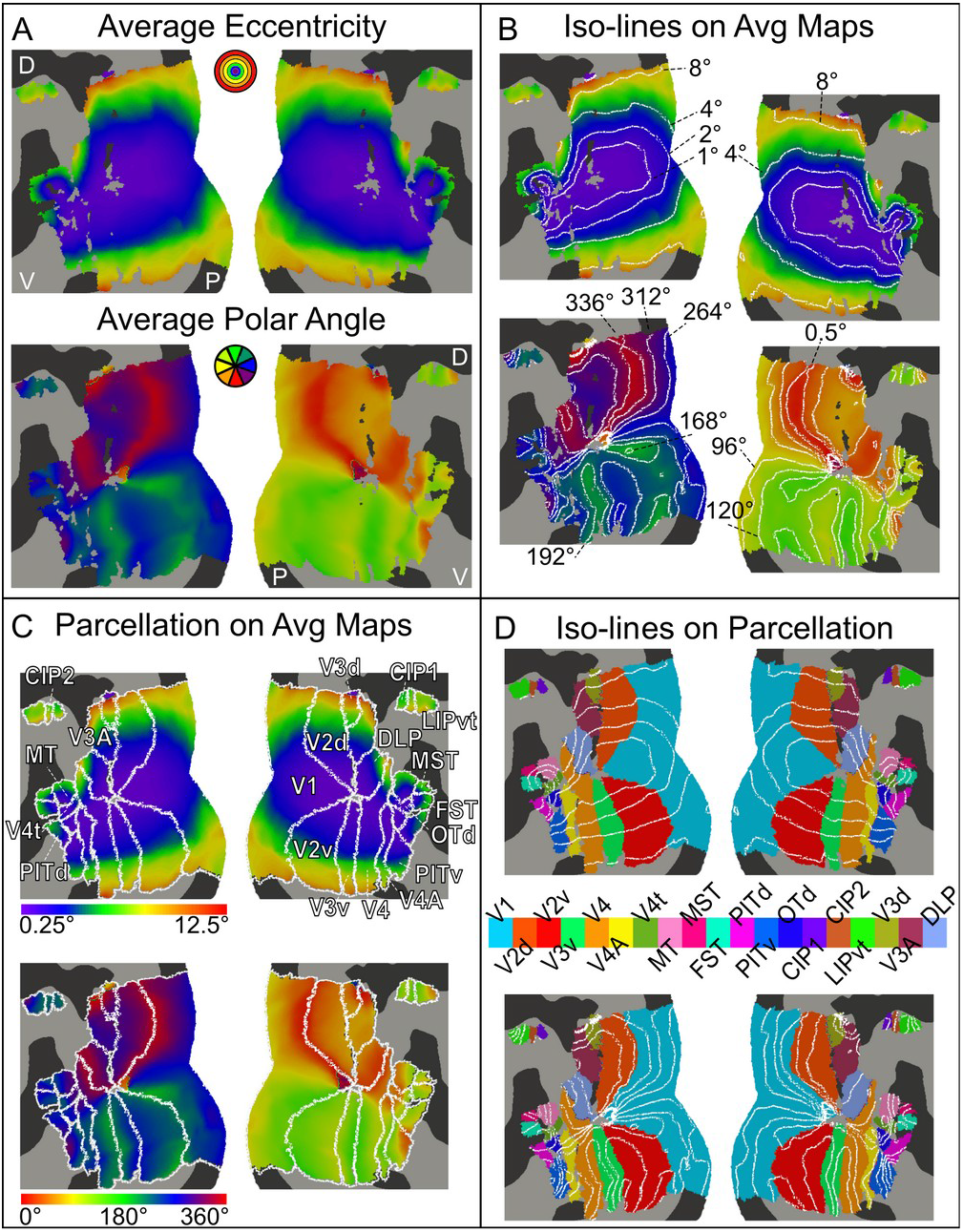
Average Maps, Iso-lines, and Parcellation (A) Average eccentricity (visual angle) and polar angle maps displayed on left and right hemisphere flatmap representations. Dorsal (D), ventral (V), and posterior (P) anatomical markers shown in white. (B) Average eccentricity (visual angle) and polar angle maps are displayed on the left and right hemisphere flatmaps, overlaid with iso-eccentricity lines and iso-polar angle lines, respectively. (C) Average eccentricity (visual angle) and polar angle maps displayed on the left and right hemisphere flatmaps overlaid with 50% probabilistic parcellation. (D) 50% Probabilistic parcellation displayed on left and right hemisphere flatmaps overlaid with iso-eccentricity lines (top) and iso-polar angle lines (bottom). The color map for individual areas is shown in the middle of the panel.

### Individual Subject Results

The primary goal of this study was to produce a probabilistic characterization of retinotopically defined boundaries of macaque visual areas. We therefore performed analyses at the individual subject level. Areal boundaries in each subject were based on thresholded (p < 0.001, uncorrected) maps of polar angle, eccentricity, and field sign, with minor adjustments made using unthresholded (raw) maps to better encompass the full extent of certain regions (see Figure S1 for example maps from M17).

For each subject, 16 visual areas were identified reliably. In three subjects (M25, M38, and M24) for whom 0.6 mm isotropic resolution data were available, up to 19 areas could be delineated. These areas do not represent the full extent of the acquired retinotopic data, but rather the subset of visual areas that have been previously defined in the literature using established retinotopic criteria. Throughout, we adhere closely to these published definitions. Although substantial evidence supports the existence of additional retinotopically defined areas, many have yet to be described and validated. We therefore excluded these regions from the present version of the probabilistic atlas.

Areal labels were projected from individual native space to the MEBRAINS template using surface-based registration^33^. For each cortical vertex and each area label, we computed the proportion of subjects that exhibited that label, yielding probabilistic ‘heatmaps’ that reflect cross-subject spatial consistency. In these maps, warmer colors indicate greater inter-individual agreement (yellow: all or nearly all subjects), whereas cooler colors indicate greater variability (red: presence in only one or two subjects). Figures 3-7 show these probabilistic maps on both the inflated surface of the cortex and flatmap reconstructions of the MEBRAINS template. Additionally, the areal boundary contours of each subject are overlaid on the flatmaps to further illustrate spatial variability across animals. In the following sections, we briefly describe the probabilistic maps for groups of anatomically neighboring visual areas.

**Figure 3:**
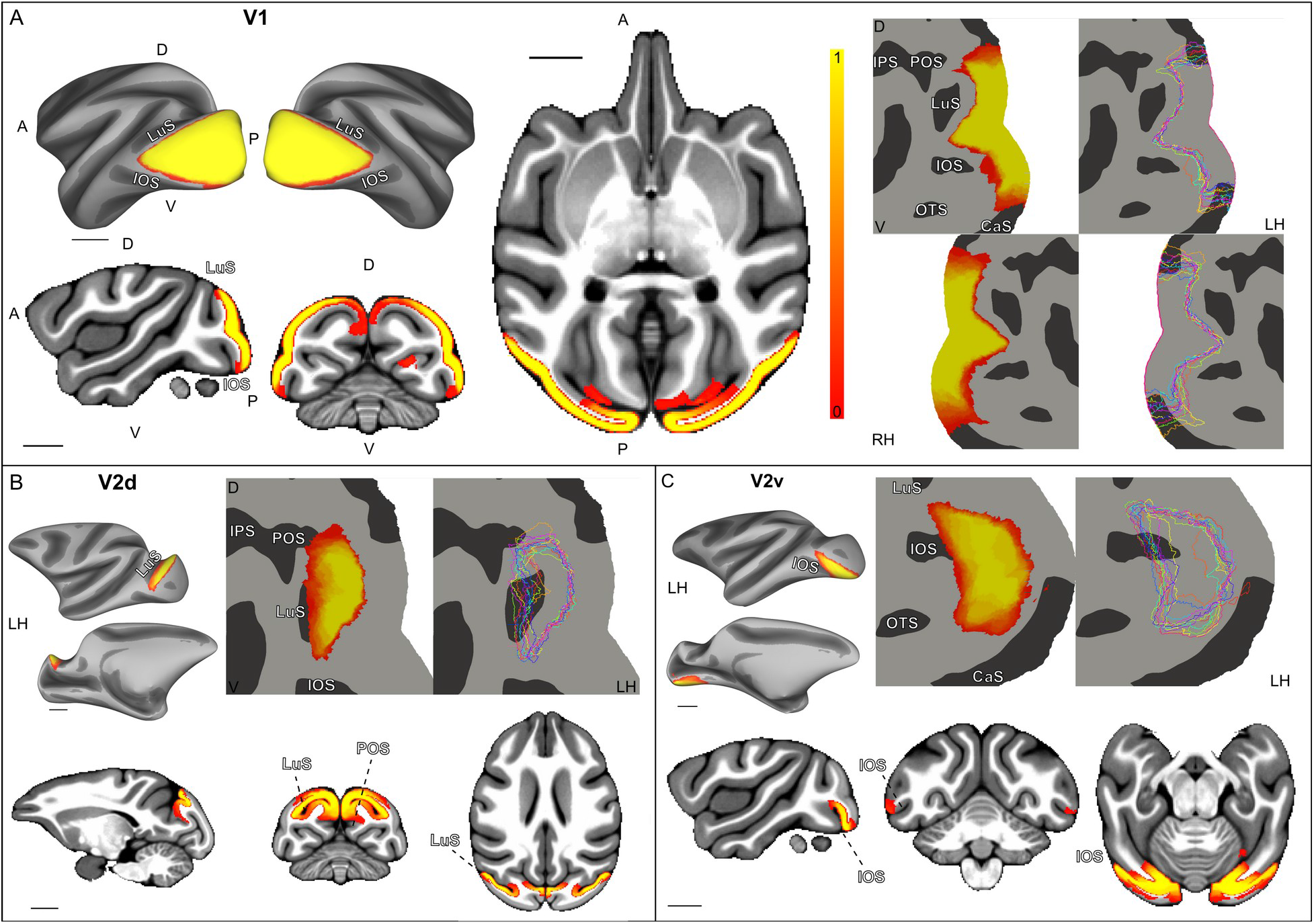
Population probabilistic heatmap and line representations of individual subject ROI definitions for (A) V1, (B) V2d, and (C) V2v. For each panel, surface, volume, and flatmap representations show first, a red/yellow (red (0) representing a single subject, i.e., no overlap, and yellow (1) representing all subjects) color overlay (heatmap) depicting the degree of overlap between the subjects’ ROI boundaries, and second, individual subject ROI lines with each color representing a different subject. Sulci names are shown in white text with black outlines. Dorsal (D), ventral (V), posterior (P), and anterior (A) anatomical markers are shown in black in panel A. Scale bars indicate 10 mm.

### V1 & V2

The boundary of V1’s visual field map is defined by the vertical meridian: the lower and upper vertical meridians are represented dorsally (V1d) and ventrally (V1v), respectively. These representations also mark the transition to the corresponding lower and upper contralateral quadrant representations of V2 (V2d and V2v), respectively (Figure 2C)^2,34–36^. Consequently, V1 comprises a single continuous cortical representation, whereas V2 is divided into dorsal (lower visual field) and ventral (upper visual field) quadrants that are separated by V1^1,35,37,38^. In both areas, the dorsal and ventral components converge at the foveal representation (see iso-eccentricity lines in Figure 2B). Anatomically, the functional boundaries representing the central 10° of the visual field closely follow the cortical folding. The transition from V1d to V2d is located at the lip of the posterior bank of the lunate sulcus (Figure 3A, B), whereas the transition from V1v to V2v lies along the posterior lip of the inferior occipital sulcus^36,39^.

These early visual areas also exhibited the highest level of inter-subject consistency. For V1, more than 90% of individual subject vertices (median across subjects) were contained within the final 50% probability map (see Table S1). Agreement was slightly lower for V2d and V2v (still exceeding 80% across hemispheres), but these areas remained among those with the greatest cross-subject consistency, as illustrated by the close correspondence of individual boundary outlines in Figure 3B and C (only the right hemisphere is shown).

The V1 and V2 boundaries also showed the strongest correspondence with previously published cortical parcellations, closely matching those reported in various maps,^21,40–42^ for review, see Vanduffel et al.,^1^. Surprisingly, however, one of the most commonly used atlases for macaque research, the CHARM atlas, places the dorsal V1/V2 slightly differently. As illustrated in Figure 8, the CHARM V1/V2 border does not align with the lip of the posterior bank of the lunate sulcus, contrary to both cytoarchitectonic features and established functional definitions of the V1/V2 boundary^43^.

### V3 & DLP &V3A

V3d is defined as a lower field representation immediately rostral to V2d, with the two areas separated by a contralateral horizontal meridian (Figure 2C)^10,44^. Slightly antero-lateral, within the lunate and parieto-occipital sulci, lies a full hemifield representation corresponding to V3A^10,39,45^. Within V3A, the lower visual field is represented dorsomedially at the annectant gyrus and the upper visual field ventrolaterally along the anterior bank of the lunate sulcus (Figure 3A), directly adjoining V2d^11,20,46^. V3d and V3a are separated by a lower vertical meridian.

Adjacent to V3A, DLP occupies a cortical sector ventrolateral to V2d and contains a complete hemifield representation centered near the foveal confluence of the early visual areas. We defined this definition of DLP^11^ based on retinotopic studies in New World monkeys, specifically the owl monkey^11,47^, although alternative schemes have proposed incorporating this region into V3A^7,48^. Along its dorso-caudal border, DLP shares an upper vertical meridian representation with V3A, whereas rostrally it borders V4 along a lower vertical meridian.

V3d, V3A, and DLP could be reliably identified in those 3 subjects in which we acquired 0.6 mm isotropic resolution fMRI data using implanted phased-array Rx coils (M25, M38, and M24)^11^. Consequently, overlap metrics were not calculated for these areas. Nevertheless, comparison of the individual areal outlines revealed a high degree of consistency across these subjects (Figures 4A, 4B, and 4D). Moreover, the population-average maps (Figure 2) permit clear delineation of V3d, V3A, and DLP, closely matching the boundaries incorporated into our final parcellation. Importantly, the existence of these 3 areas was confirmed in an additional cohort of 8 subjects who were scanned at low resolution^11^. Relative to previous parcellations, our scheme subdivides what has often been described as a single dorsal V3d region into V3d, (part of) V3A, and DLP, while preserving the overall external boundaries of this complex.

**Figure 4:**
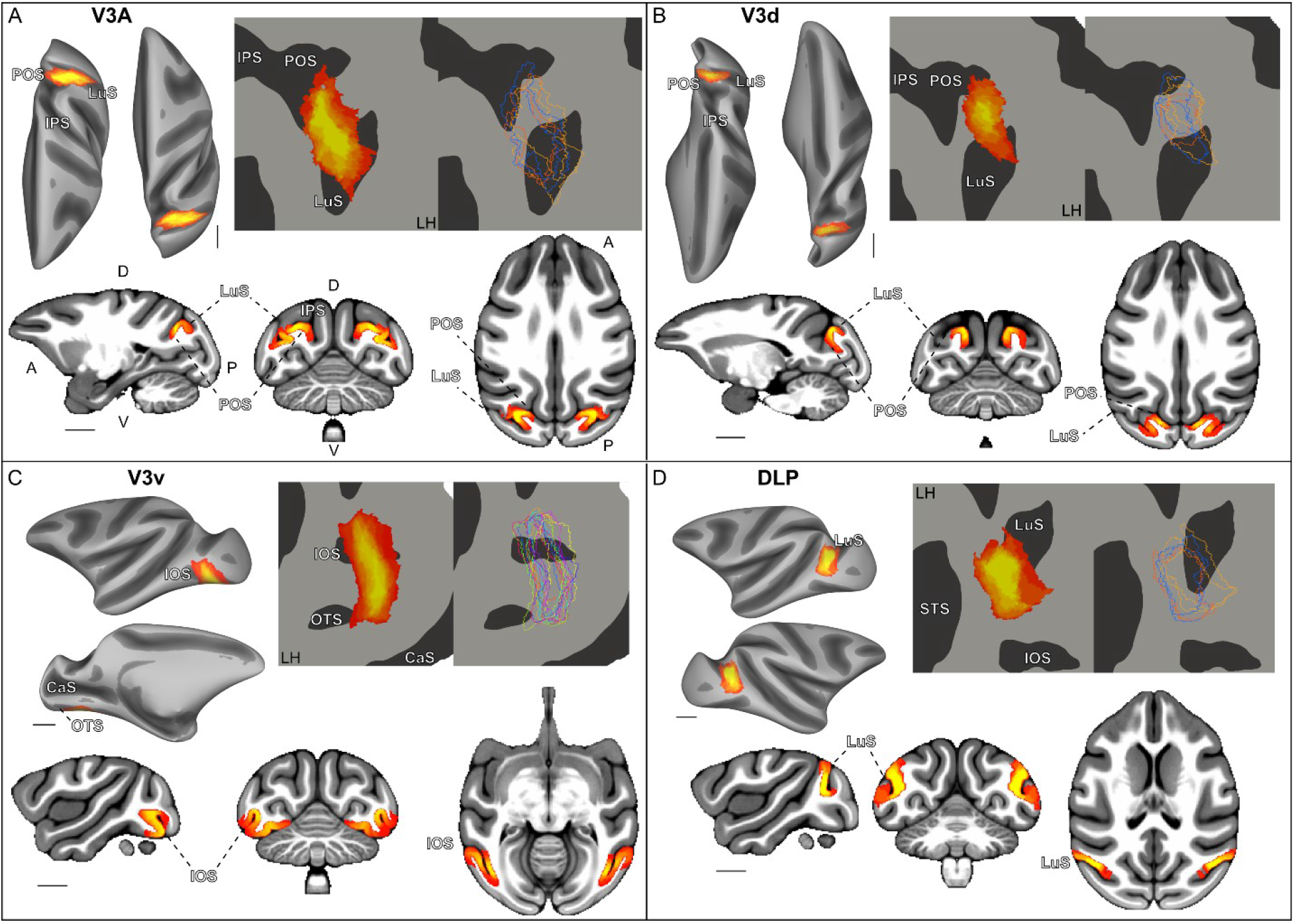
Population probabilistic heatmap and line representations of individual subject ROI definitions for (A) V3A, (B) V3d, (C) V3v, and (D) DLP. For each panel, surface, volume, and flatmap representations show first, a red/yellow (red (0) representing a single subject, i.e., no overlap and yellow (1) representing all subjects) color overlay (heatmap) depicting the degree of overlap between the subjects’ ROI boundaries, and second, individual subject ROI lines with each color representing a different subject. Sulci names are shown in white text with black outlines. Dorsal (D), ventral (V), posterior (P), and anterior (A) anatomical markers are shown in black in panel A. Scale bars indicate 10 mm.

The upper field representation immediately rostral to V2v corresponds to V3v, with the areas separated by a horizontal meridian. V3v is separated from V4 by an upper vertical meridian, consistent with previous electrophysiological and functional mapping studies^7,49–51^. Although several previous parcellations have incorporated part of this territory into V2 (see Figure 8), our delineation of V3v is consistent with the organization proposed by Markov et al^40^.

### V4 and V4A

V4 stretches between the lunate sulcus (LuS) and STS, and the inferior occipital sulcus (IOS) and the STS, covering parts of the prelunate gyrus and the inferior temporal gyrus (see Figure 5A)^46,52,53^. Ventrally, its rostral boundary is defined by a horizontal meridian separating V4 from V4A^11,13^. The latter area contains an entire contralateral hemifield representation, with the lower and upper quadrant representations positioned rostral to the dorsal and ventral portions of V4, respectively^13,29^.

**Figure 5:**
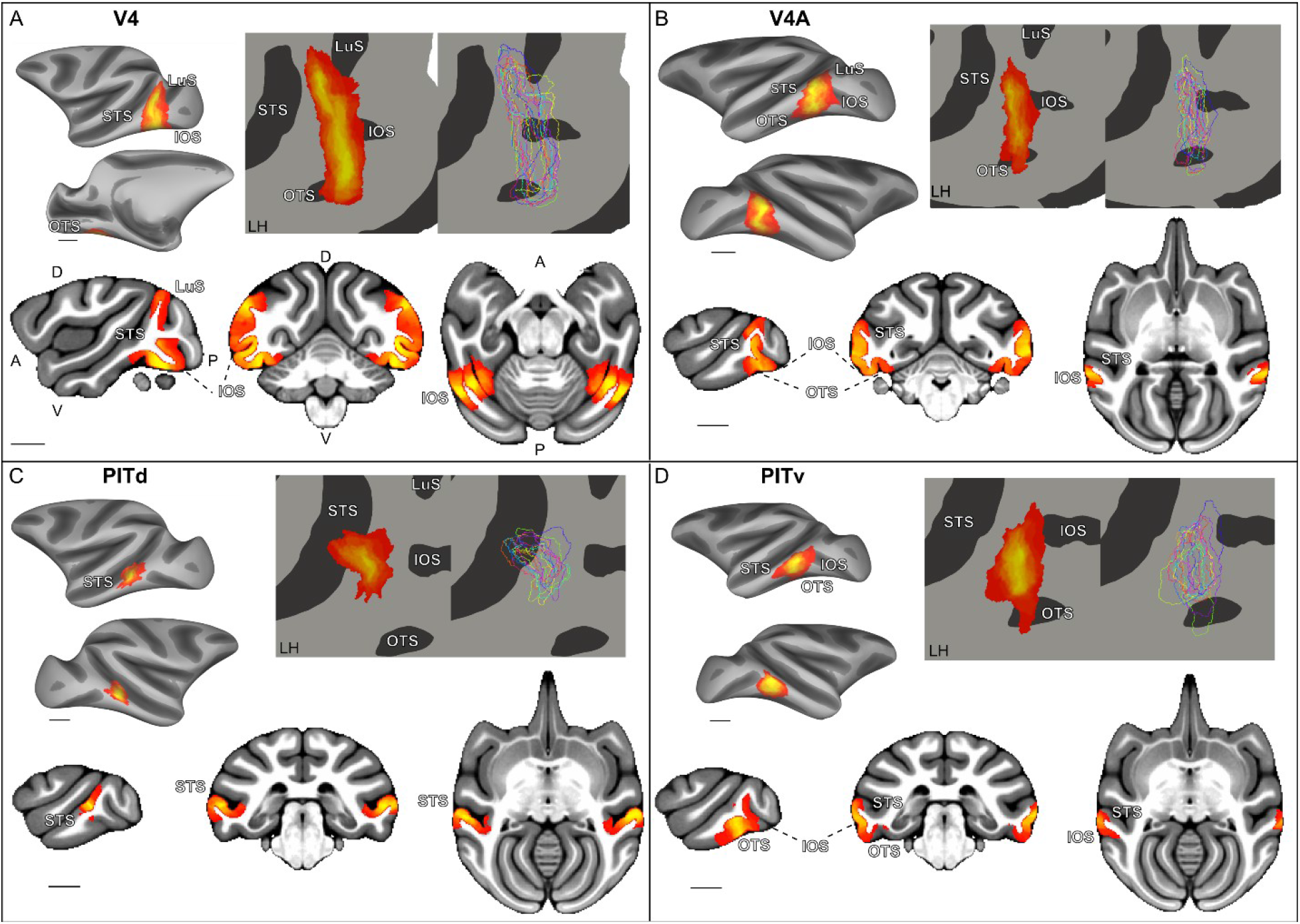
Population probabilistic heatmap and line representations of individual subject ROI definitions for (A) V4, (B) V4A, (C) PITd, and (D) PITv. For each panel, surface, volume, and flatmap representations show first, a red/yellow (red (0) representing a single subject, i.e., no overlap and yellow (1) representing all subjects) color overlay (heatmap) depicting the degree of overlap between the subjects’ ROI boundaries, and second, individual subject ROI lines with each color representing a different subject. Sulci names are shown in white text with black outlines. Dorsal (D), ventral (V), posterior (P), and anterior (A) anatomical markers are shown in black in panel A. Scale bars indicate 10 mm.

Although V4A was originally described more than five decades ago^54^, several widely used parcellation schemes continue to incorporate it into V4. This convention is difficult to reconcile with evidence that V4A contains an independent contralateral hemifield representation^55^ and distinct patterns of cortical connectivity relative to V4^56^. In our dataset, V4 showed a high degree of spatial consistency across subjects, with >70% vertex overlap. V4A, however, was slightly more variable, exhibiting ∼ 50% vertex overlap across subjects. However, as illustrated in the upper right panel of Figure 5B, this lower overlap was primarily driven by a few outlier subjects rather than widespread ambiguity in the location of the area.

### MT Cluster

MT, MST(v), FST, and V4t each contain a complete contralateral hemifield representation and lie on the posterior bank of the dorsal sector of the STS (Figures 6 A-B and 7A)^9,57–60^. These motion-sensitive areas form a ‘cluster’ organized around a shared foveal representation (Figure 2B)^13,57^, with MT positioned anterior to the dorsal portion of V4 and medial to V4t, and MSTv and FST progressively more anterior^61–64^. This cluster can be identified by the ‘cross-like’ configuration formed by alternating lower vertical, horizontal and upper vertical meridian representations and separated from early visual areas (V1– V4) by an eccentricity “ridge”^13^. Within this cluster, spatial consistency varied across areas. MT is by far the most reproducible, with > 65 % vertex overlap across subjects (Table S1). In contrast, MSTv, FST, and V4t each showed lower overlap values (<60%).

**Figure 6:**
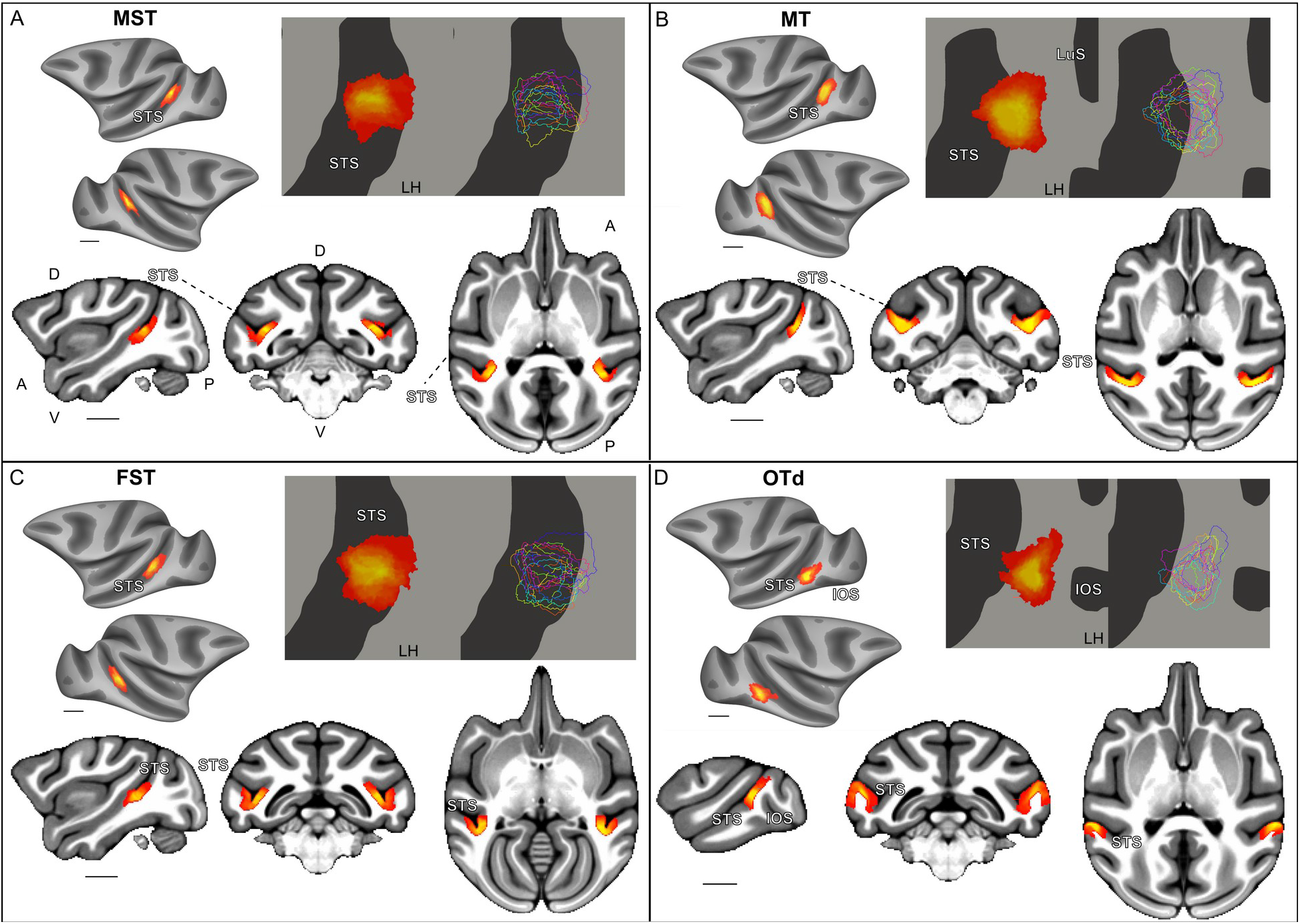
Population probabilistic heatmap and line representations of individual subject ROI definitions for (A) MST, (B) MT, (C) FST, and (D) OTd. For each panel, surface, volume, and flatmap representations show first, a red/yellow (red (0) representing a single subject, i.e., no overlap and yellow (1) representing all subjects) color overlay (heatmap) depicting the degree of overlap between the subjects’ ROI boundaries, and second, individual subject ROI lines with each color representing a different subject. Sulci names are shown in white text with black outlines. Dorsal (D), ventral (V), posterior (P), and anterior (A) anatomical markers are shown in black in panel A. Scale bars indicate 10 mm.

Comparison of our parcellation with the CHARM atlas reveals good overall agreement for several areas, particularly MT (Figure 8, probabilistic parcellation in pink, CHARM’s borders labeled as 7). A similar correspondence is observed for FST (green, number 9), but not for MST or V4t. The region labeled as MST in CHARM (which does not distinguish dorsal and ventral subdivisions) is more dorsal and anterior than the region we identify as MSTv and overlaps primarily with the CHARM-defined FST. This discrepancy may partially reflect differences in the range of eccentricities sampled in our retinotopic mapping, as MST neurons are known to represent a much larger eccentricity range than was covered by our stimuli^62^. An alternative possibility is that the CHARM parcellation primarily captures MSTd while incorporating MSTv into FST, thereby obscuring the distinction between these two areas.

### Inferotemporal Cluster

Ventral to this ‘MT cluster’, but still along the posterior bank of the STS, V4A, PITd, PITv, and OTd form another cluster (Figures 5B-D and 6D), with each area also containing a complete hemifield representation^8,13,29^. Within this cluster, OTd, PITd, and PITv lie anterior to V4A, with OTd separated from both PIT compartments by a vertical meridian representation. The most anterior extent of PITd/PITv is defined by another vertical meridian. Interestingly, PITv and PITd exhibited a higher degree of overlap across subjects in the right than in the left hemisphere (Table S1). Notably, as illustrated by the overlap of our parcellation (Figure 8), PITv, PITd, and OTd encompass territory that has frequently been described as TEO or VOT in previous parcellation studies^42,65,66^.

### Parietal areas CIP1, CIP2 and LIPvt

In the parietal cortex near the transition from the parieto-occipital sulcus (POS) and the intraparietal sulcus (IPS), we identified a complete hemifield representation in LIPvt and two half-hemifield representations in CIP1 and CIP2^67^. CIP1 is separated from V3A by an upper vertical meridian representation^10^. A horizontal meridian, on the other hand, marks the transition from CIP1 to CIP2, which has a mirror-reversed half-hemifield organization relative to CIP1 (Figure 2). The upper vertical meridian rostral to CIP2 marks the border with LIPvt^68–70^. These areas are located on the posterior bank of the IPS (Figure 7B-D). CIP1 and CIP2 showed comparatively lower spatial consistency across subjects, with less than 50% of overlapping vertices, making them the most heterogeneous regions considered in this study.

**Figure 7:**
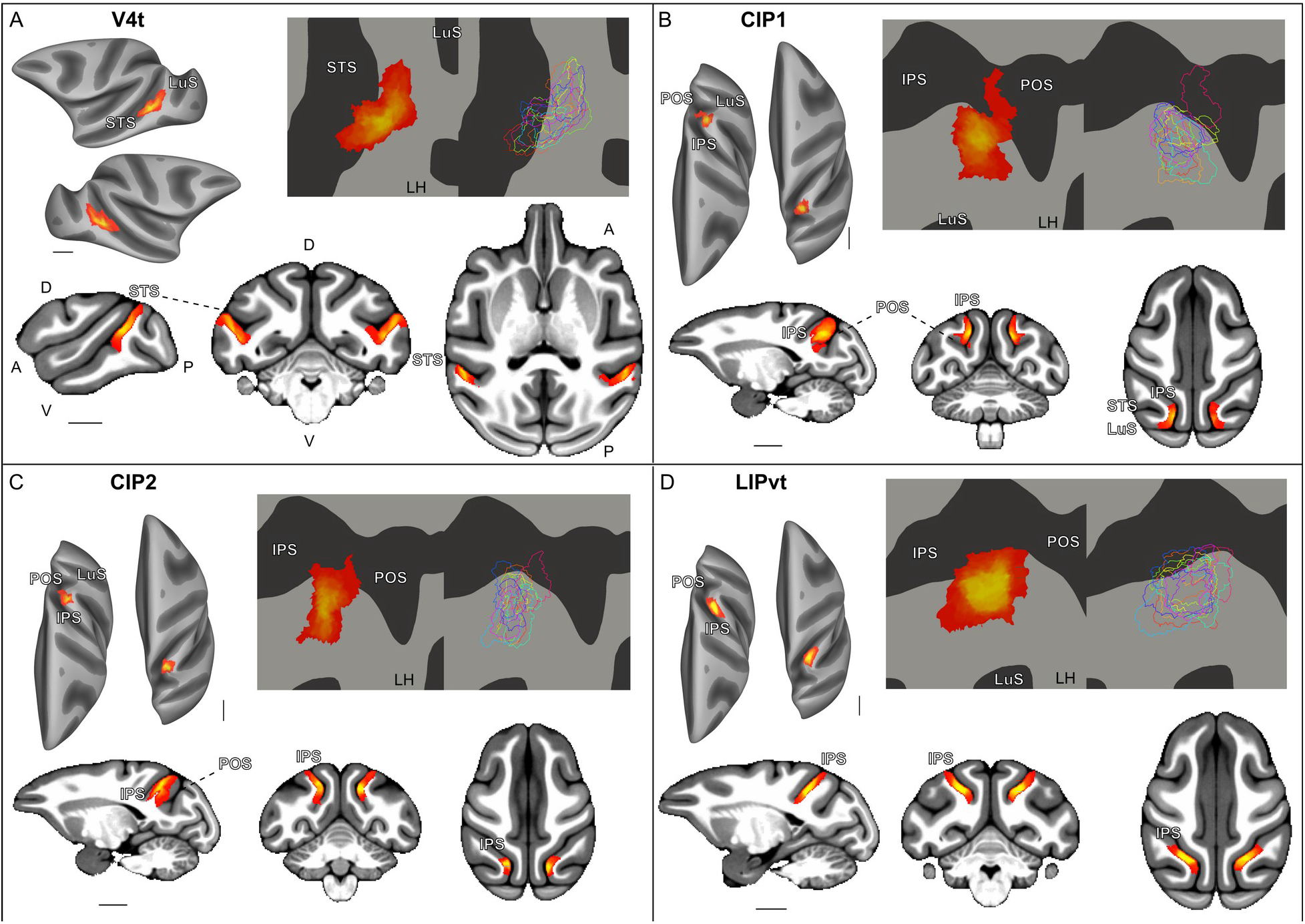
Population probabilistic heatmap and line representations of individual subject ROI definitions for (A) V4t, (B) CIP1, (C) CIP2, and (D) LIPvt. For each panel, surface, volume, and flatmap representations show first, a red/yellow (red (0) representing a single subject, i.e., no overlap and yellow (1) representing all subjects) color overlay (heatmap) depicting the degree of overlap between the subjects’ ROI boundaries, and second, individual subject ROI lines with each color representing a different subject. Sulci names are shown in white text with black outlines. Dorsal (D), ventral (V), posterior (P), and anterior (A) anatomical markers are shown in black in panel A. Scale bars indicate 10 mm.

Here too, we diverge most heavily from previously published parcellations. However, our CIP1/2 corresponds exactly with those described in Lewis and Van Essen^42^, and our LIPvt does fall within the bounds of what others have described as LIP(v)^40–42^. When we compare to CHARM, these three areas follow the border of LIPd/v, which may indicate that we are simply facing a coverage problem and that these three areas represent a different subdivision parcellation of LIP.

### Inter-subject Consistency

Across the 13 subjects, inter-subject spatial consistency followed a caudo-rostral gradient across the visual cortex. Early visual areas, particularly V1 and V2, exhibited the highest correspondence with the final parcellation, whereas progressively greater variability was observed in more anterior extrastriate and posterior parietal areas (Table S1).

This trend is also evident in the probabilistic maps shown in Figure S2. Early visual areas exhibit sharply defined regions with high overlap, while the probability distributions become progressively broader in higher-order visual areas. Although individual areas within the MT cluster remain distinguishable, their boundaries show greater spatial variability across subjects than those of early visual cortex. Because these maps represent the full spatial probability distribution for each area, some broadening is expected; nevertheless, the progressive increase in variability from posterior to anterior cortex is readily apparent.

**Figure 8:**
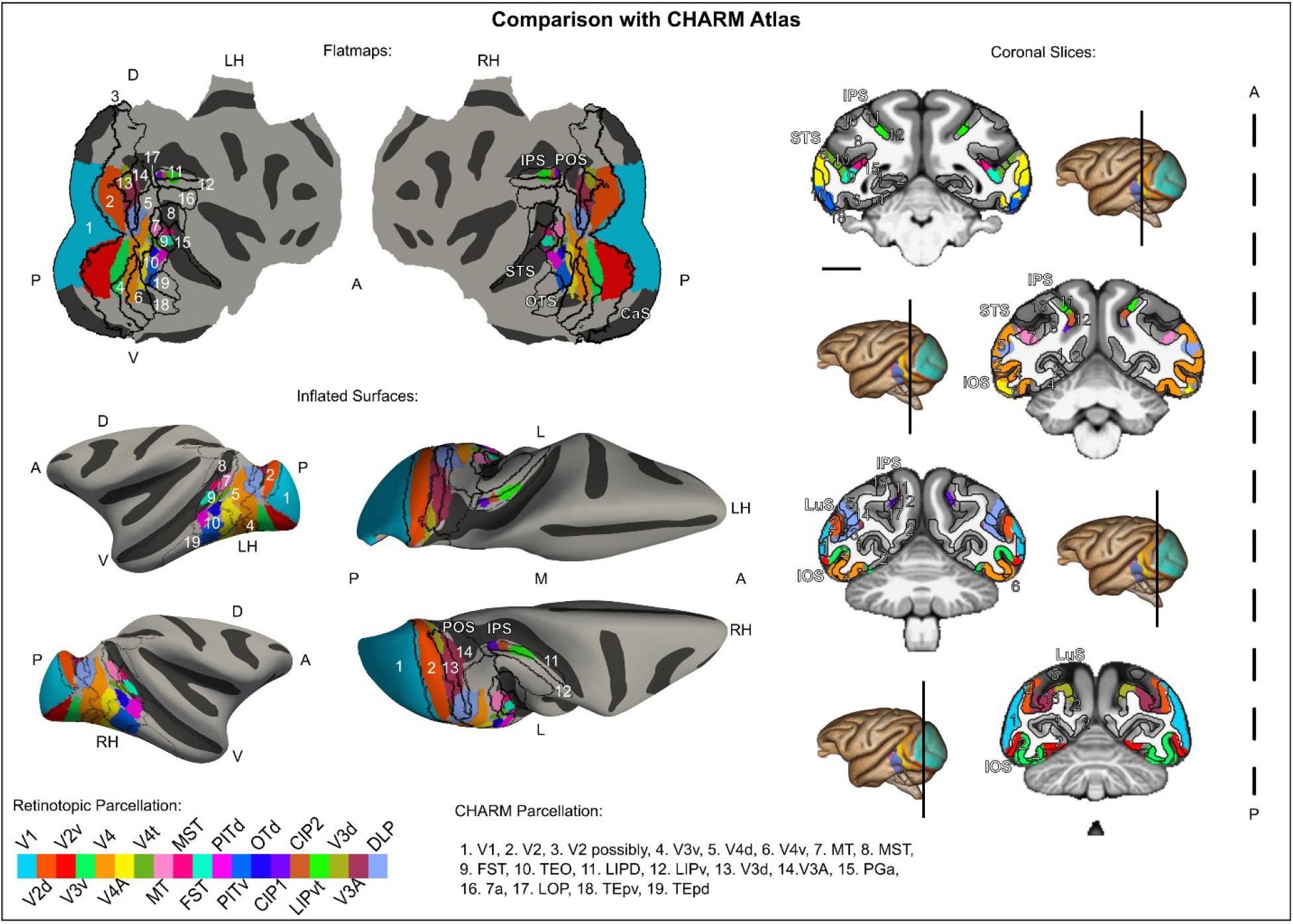
Comparison of Retinotopic Parcellation with CHARM Atlas. The MEBRAINS template is shown in three formats: flatmaps (left and right hemispheres), inflated surfaces viewed from the lateral extent (left and right hemispheres) and from above (left and right hemispheres), and four cortical slices. In each view, the retinotopic parcellations are shown with the same color mapping as in Figure 2. Overlaid in white are the corresponding labels from the CHARM atlas. Scale bar indicates 10 mm.

## Discussion

To our knowledge, this dataset represents the largest phase-encoded retinotopic fMRI resource acquired in macaques to date. Using this data, we delineate 19 retinotopically organized visual areas and provide both individual-subject areal boundaries and probabilistic maps that characterize the spatial distribution of each area across animals. All maps are registered to the MEBRAINS macaque anatomical template^32^ and are openly available through the EBRAINS data repository (URL *will be released upon acceptance*).

A key feature of this resource is the inclusion of quantitative measures of inter-individual spatial overlap. These analyses reveal a gradient of anatomical consistency across the visual hierarchy: early visual areas, such as V1 and V2, show relatively high spatial consistency across subjects, whereas higher-order regions, including V4A, CIP1, and CIP2, exhibit substantially larger variability. The basis of this increased variability remains unclear. It may reflect anatomical factors, such as greater inter-individual differences in cortical folding and areal morphology in higher-order cortex, functional factors associated with increasingly complex computations and more flexible organizational principles, or an interaction between these anatomical and functional influences. This resource provides a solid quantitative framework for investigating these questions in future studies.

### Challenges to current models of macaque visual cortex organization

The probabilistic parcellation presented here highlights several discrepancies with widely accepted models of macaque visual cortex organization. These differences emerge early in the visual hierarchy, particularly in the cortex immediately anterior to dorsal V2. In contrast to conventional parcellations (Jeffs et al., and Gattass et al., ^48,71^), our data support the presence of three distinct areas (V3d, V3A, and DLP), an organization that more closely resembles that reported in New World monkeys^11,72^. Notably, this interpretation is also consistent with the original delineation of V3A by Van Essen and Zeki^46^.

Our data also support recognizing V4A as a distinct visual area immediately rostral to ventral V4. Although V4A was originally described more than five decades ago^54^, it is absent from many commonly used macaque cortical parcellations, where it is instead incorporated into ventral V4 or TEO^21,40–42^. However, previous publications have demonstrated that this region contains a complete representation of the contralateral hemifield^13,29^, a finding that is robustly replicated in the present dataset. Together, these observations argue that V4A is more appropriately considered a distinct retinotopic area rather than a subdivision of neighboring cortex.

Overall, our delineation of the MT cluster is largely consistent with previous parcellations, with broad agreement on the locations of MT, MST, FST, and V4t. A limitation of the present dataset is the limited eccentricity range sampled (up to 12.5°), which did not capture the full retinotopic extent of these areas. This limitation may also influence our definitions of CIP1 and CIP2^67^ and LIPvt^73^. Despite these constraints, our findings support a robust retinotopic organization across the MT cluster. It should be noted that the largest discrepancy between our parcellation and the CHARM atlas^74^ concerns MST. The CHARM delineation of MST as a single undivided area can be traced back to the original D99 atlas^41^, which provides a digital 3D reconstruction of the Saleem and Logothetis atlas ^22^. In the original atlas, MST subdivisions were informed by functional studies but were primarily delineated based on differences in the distribution of parvalbumin and SMI-32 staining in a single subject^22^. Notably, among the functional studies used to guide these definitions, the locations of MT, FST, and V4t in our parcellation correspond closely to those defined by^64,75^. Furthermore, our MSTv subdivision corresponds well to their DMZ, a “densely myelinated zone” located predominantly within MST along the upper bank of STS.

A more substantive difference from previous parcellations concerns the cortical territory traditionally assigned to TEO. We suggest that this discrepancy primarily reflects differences in parcellation criteria rather than fundamental disagreement about cortical organization. Whereas anatomical approaches have treated this region as a single area, our retinotopic analyses indicate that the anterior bank of the occipitotemporal sulcus extending into the inferior temporal gyrus is more accurately subdivided into V4A, PITd, PITv, and OTd. These regions may therefore be viewed as retinotopically distinct subdivisions within the broader anatomical territory mostly encompassing TEO.

### Limitations and Potential Sources of Error

Differences between our parcellation and the most commonly used macaque atlases are discussed in greater detail by Vanduffel and Zhu (2025)^1^. One potential source of discrepancy is the limited retinotopic window sampled in the present dataset (up to 12.5° eccentricity). This constraint primarily reflects a technical limitation of retinotopic fMRI at 3T, although larger eccentricity ranges have been achieved using specialized experimental adaptations^10^ and in more recent studies extending to 40° eccentricity^12^. Nevertheless, because a substantial proportion of macaque visual cortex represents the central 12.5° of the visual field, our maps capture the majority of each area’s retinotopic organization. The principal consequence is that the boundaries of the most anterior visual areas may remain incompletely defined. Representing these areas probabilistically allows this uncertainty to be captured explicitly rather than imposing deterministic borders.

More generally, it should be appreciated that uncertainty in cortical parcellation is not unique to fMRI. Electrophysiological maps are assembled from sequential recordings with finite spatial sampling and require reconstruction of recording locations across many experiments and subjects. Likewise, cytoarchitectonic, receptor-architectonic, and connectivity-based parcellations each rely on distinct criteria and are subject to their own methodological limitations and inter-subject variability. Functional imaging offers complementary advantages by enabling dense, whole-brain measurements of retinotopic organization within individual animals, but it also introduces modality-specific limitations, including the restricted eccentricity range described above.

It is important to note that reliance on a single subject represents a major limitation of many traditional and commonly used atlases. Our results demonstrate that single-subject parcellations, including the CHARM/D99 atlas, can differ substantially from population-based functional organization. For example, in the case of the precise V1/V2 boundary discussed above, the discrepancy is inconsistent with established anatomical and functional definitions of these areas.

Another source of uncertainty arises from inter-subject registration. Transforming data from 13 individual native surfaces to a common template inevitably introduces some degree of spatial ambiguity. We expect this to be substantially reduced by surface-based registration, which better preserves cortical topology than volumetric approaches^76^. Moreover, representing areal boundaries probabilistically, rather than as fixed deterministic borders, naturally accommodates residual inter-subject variability and registration uncertainty, providing a more faithful representation of confidence in areal localization.

## Conclusion

This dataset provides a state-of-the-art probabilistic retinotopic atlas of the macaque visual cortex that serves as a resource for interpreting, designing, and guiding future neuroimaging, electrophysiological, and causal intervention studies in macaques. In addition to refining the delineation of established visual areas, it provides evidence supporting the inclusion of previously overlooked or underrepresented areas, such as V4A, in contemporary macaque visual atlases. By explicitly quantifying inter-subject variability through overlap metrics and probabilistic maps, the atlas provides a principled framework for anatomical localization while representing uncertainty in areal boundaries and room for future adaptations. More broadly, this resource complements the growing ecosystem of probabilistic brain atlases and contributes to increasingly reproducible and quantitative analyses in systems neuroscience.

Finally, although the present parcellation should not be viewed as a definitive “ground truth” map of the macaque visual cortex, its integration within MEBRAINS, alongside cytoarchitectonic, receptor-architectonic, and connectivity-based datasets, provides a framework for systematically and objectively comparing complementary parcellation schemes^77^. Such integration will facilitate the convergence of multiple lines of evidence toward an increasingly comprehensive and biologically grounded atlas of the macaque brain.

## Acknowledgments

The authors thank J. Arsenault, X. Li, N. Caspari, A. Gerits, P. Balan, and S. Bensemmane for data collection as well as C. Fransen, I. Puttemans, A. Hermans, S. Verstraeten, W. Depuydt, and M. De Paep for technical and administrative support. This project received financial support from the following: KU Leuven: C14/21/111; Fonds Wetenschappelijk Onderzoek-Vlaanderen (FWO-Flanders): G0E0520N, G0C1920N; European Union’s Horizon 2020 Framework Programme for Research and Innovation: Human Brain Project SGA3; EBRAINS 2.0: ZL37682100-103

## Author Contributions

Conceptualization: Q.Z. and W.V.; methodology: Q.Z. and W.V.; formal analysis: N.E.F. and Q.Z.; visualization and data curation: N.E.F.; writing (original draft): N.E.F. and W.V.; writing (review & editing): N.E.F., Q.Z., and W.V.; funding acquisition: W.V.; supervision: W.V.

## Supplemental information

Document S1. Table S1, Figure S1-2

Video S1. Representation of parcellation in volume and coronal slices, related to Figure 2.

## Abbreviations & Definition Sources

V1/V2: Visual areas 1-2 ^1,2,34–38,43^
V3: Visual area 3
V3d: Dorsal area V3 ^10,44^
V3A: accessory area V3 ^10,11,20,39,45,46^
DLP: Dorsolateral posterior area ^11,47^
V3v: Ventral area V3 ^7,49–51^
V4: Visual area 4 ^46,52,53^

**MT Cluster^9,57–64^**

V4t: Transitional area V4
MT: Middle temporal area
FST: Fundus of superior temporal area
MSTv: Medial superior temporal area, ventral part

**Inferotemporal Cluster ^8,13,29^**

OTd: Dorsal occipitotemporal area
PITd: Dorsal posterior inferotemporal area
PITv: Ventral posterior inferotemporal areas
V4A: anterior area V4 ^54,55^

**Parietal Areas ^10,67–70^**

CIP1: Caudal intraparietal area 1
CIP2: Caudal intraparietal area 2
LIPvt: Visuotopic lateral intraparietal area
fMRI: Functional magnetic resonance imaging

## Methods

### 1. Experimental Model and Study Participant Details

#### Subjects

Thirteen subjects (*Macaca mulatta*) were used in this study. Animal care and experimental procedures were performed in accordance with the National Institute of Health’s Guide for the Care and Use of Laboratory Animals, the European legislation (Directive 2010/63/EU), and were approved by the Animal Ethics Committee of KU Leuven. Housing and handling were in accordance with the Weatherall report. All animals were group-housed in cages sized 16-32 m^3^, which encourages natural social interactions and locomotor behavior. The environment was enriched by foraging devices and toys. The health of the subjects was monitored and maintained daily by trained technical staff, veterinary staff, and experimenters, all trained to work with NHPs. For each subject, MR-compatible headposts were surgically implanted and secured with ceramic screws and dental cement as previously described in ^78^. NHPs had unrestricted access to food and daily access to restricted volumes of fruits and water; during experiments, NHPs had unlimited access to fluids as part of the performance-based protocol. NHPs had unrestricted access to food and daily access to restricted volumes of fruits and water; during experiments, NHPs had unlimited access to fluids as part of the performance-based protocol.

### 2. Method Details

#### Experimental Controls

For all scanning sessions, subjects were seated in a head-fixed sphinx position inside a custom-built plastic primate chair, which was positioned in the center of the scanner bore. During each run, subjects were trained to maintain visual fixation within a 2° × 2° virtual window in the center of the screen. Additionally, subjects were trained to keep their hands positioned within a small response box in front of the chair ^79^. Hand positions were continuously checked with two pairs of optical fibers. Rewards (in the form of liquid juice droplets) were contingent upon correct eye fixation behavior and having both hands inside this response box. This procedure significantly reduced body motion, which resulted in a higher quality of the EPI’s compared to controlling solely for eye fixation behavior.

#### (f)MRI acquisition

For each subject, 8–11 mg/kg of one of three iron-based contrast agents (monocrystalline iron oxide nanoparticle (Molday ION, BioPAL), Sinerem (Guerbet), or Feraheme (AMAG Pharmaceuticals)) was injected via the femoral/saphenous vein to improve the contrast-to-noise ratio (CNR) and to avoid the contribution of superficial draining veins ^78,80^. To reduce the risk of iron accumulation, 1-1.5 g/day deferoxamine mesylate (Desferal, Novartis; intramuscular injection) was administered immediately after the scan, for a period of 3–7 days, until serum iron and ferritin levels returned approximately to normal ranges (confirmed by blood test).

All scans were acquired using a 3T Siemens TIM-TRIO, except for the scans of M38, which were acquired using a 3T Siemens Prisma Fit scanner. For M25, M35, and M24, data were acquired with an AC88 gradient insert. Only scan sessions with excellent behavioral performance (above 90% during the total scan duration) were included in the statistical analysis. The functional images were acquired at different resolutions depending on the type of coils used (Table 1).

**Table 1.** Subject Details for Functional Scans.

| Subject | Sex | Weight (kg) | Coil Type | Runs (> 90%) | Voxel Size (mm) | TR (ms) | TE (ms) | Flip angle (°) | Dimensions |
| --- | --- | --- | --- | --- | --- | --- | --- | --- | --- |
| M19 | M | 4.25 | External (16 ch) | 91 | 1 | 2000 | 19 | 75 | 84 x 84 x 50 x 133 |
| M25 | F | 4.4 | Implanted (8 Ch) | 29 | 0.6 | 3000 | 21 | 90 | 140 x 140 x 48 x 133 |
| M38 | M | 5.9 | Implanted (8 Ch) | 56 | 0.6 | 3000 | 22 | 90 | 140 x 140 x 74 x 135 |
| M17 | M | 3.45 | External (8 Ch) | 63 | 1 | 2000 | 19 | 75 | 84 x 84 x 50 x 133 |
| M13 | F | 4.05 | External (16 ch) | 50 | 1 | 2000 | 19 | 75 | 84 x 84 x 50 x 133 |
| M35 | M | 6.65 | Implanted (8 Ch) | 72 | 1 | 2000 | 19 | 84 | 84 x 84 x 44 x 133 |
| M14 | M | 4.25 | External (8 Ch) | 64 | 1 | 2000 | 19 | 75 | 84 x 84 x 50 x 133 |
| M18 | M | 5.15 | External (8 Ch) | 86 | 1 | 2000 | 19 | 75 | 84 x 84 x 50 x 133 |
| M24 | F | 4.1 | Implanted (8 Ch) | 156 | 0.6 | 3000 | 15 | 90 | 140 x 140 x 67 x 133 |
| M33 | M | 4.35 | External (8 Ch) | 75 | 1 | 2000 | 18 | 76 | 84 x 84 x 50 x 133 |
| M32 | M | 6.2 | Implanted (6 Ch) | 65 | 1.25 | 2000 | 19 | 75 | 84 x 84 x 50 x 133 |
| M28 | F | 5.05 | Implanted (6 Ch) | 6 | 1 | 2000 | 18 | 76 | 84 x 84 x 50 x 133 |
| M22 | M | 7.4 | External (8Ch) | 63 | 1 | 2000 | 18 | 76 | 84 x 84 x 50 x 133 |

For each subject ( $n = 13$ ), their sex, weight at the time of scanning, coil type, and details of experimental scanning (number of runs with > 90% fixation, voxel size, TR, TE, TI, flip angle, and dimensions) are provided.

##### Implanted Coils

For 6 subjects (M25/M38/M35/M24/M32/M28), 6- or 8-channel phased-array receive coils were embedded in an MRI-compatible cement implant on the skull according to ^80^. For subjects M25, M38, and M24 (previously described in Zhu et al.,^11^, we acquired sub-millimeter (0.6 mm isotropic voxel size) functional measurements. For M24 and 25, data were acquired with an AC88 gradient insert in the TIM-TRIO scanner; only a limited number of slices could be scanned at a TR of 3000ms; as such, data from the top and bottom parts of the brain of these two subjects were acquired in different sessions. For M38, it was possible to use a simultaneous multiple slice (SMS) EPI sequence to cover the whole brain (in the Prisma Fit scanner).

In addition to the EPI images, several T1 weighted (T1-w) 3D gradient echo (GRE) images (0.6 mm isotropic voxel size, TR = 6 ms, TE = 2.63 ms, α = 11°, matrix size = 160 × 160 × 96) were acquired in every fMRI session, to use as an intermediate image for the registration between functional and high-resolution anatomical images of the same subject.

##### External Coils

For the remaining 7 subjects (M19/M17/M13/M14/M18/M33/M22), custom-made external coils were used. A local single-loop transmit coil was positioned over the head, along with either an external 16- or 8-channel phased-array receive coil tightly secured on the head. Each functional scan consisted of T2* weighted echo-planar whole-brain images (EPI).

To enhance the registration between the EPI template images and the anatomical reference images (MPRAGE), 3D gradient-echo (GRE) images were acquired at the beginning of each scanning day for each monkey. Distortion correction was performed based on EPI images to the same-session GRE image warping only in the phase-encoding direction. A gradient-recalled echo field map (GRE-FM) was acquired for subjects M22 and M33 at the same EPI voxel resolution. The field maps consisted of two images acquired with different echo times (1 mm isotropic voxel size, TR = 6 ms, TE1 = 5.2 ms, TE2 = 7.66 ms; α = 11°; matrix size 84 ×84 x 50) and were used to correct for EPI distortions resulting from magnetic field inhomogeneity.

#### Reference Anatomical Images

Prior to the experiments, we acquired 10 – 15, high-resolution (0.4 mm isotropic voxel size) T1w images when the subjects were under ketamine-xylazine or ketamine-medetomidine anesthesia. For full details, including TR and TE time, see Table 2.

**Table 2.** Subject Details for High-Resolution Anatomical Scans.

| Subject | Scanner | Voxel Size (mm) | TR (ms) | TE (ms) | TI (ms) | Flip angle (°) | Dimensions |
| --- | --- | --- | --- | --- | --- | --- | --- |
| M19 | Trio | 0.4 | 2500 | 4.35 | 850 | 9 | 320 x 260 x 208 |
| M25 | Trio | 0.4 | 2500 | 4.35 | 850 | 9 | 320 x 260 x 208 |
| M38 | Prisma | 0.4 | 2700 | 3.5 | 882 | 9 | 320 x 260 x 208 |
| M17 | Trio | 0.4 | 2700 | 3.35 | 850 | 9 | 320 x 260 x 208 |
| M13 | Trio | 0.4 | 2500 | 4.35 | 850 | 9 | 320 x 260 x 208 |
| M35 | Trio | 0.4 | 2700 | 3.35 | 850 | 9 | 320 x 260 x 208 |
| M14 | Trio | 0.4 | 2200 | 2.52 | 900 | 9 | 320 x 260 x 208 |
| M18 | Trio | 0.4 | 2200 | 2.52 | 900 | 9 | 320 x 260 x 208 |
| M24 | Trio | 0.4 | 2500 | 4.35 | 850 | 9 | 320 x 260 x 208 |
| M33 | Trio | 0.4 | 2700 | 3.35 | 850 | 9 | 320 x 260 x 208 |
| M32 | Trio | 0.4 | 2700 | 3.35 | 850 | 9 | 320 x 260 x 208 |
| M28 | Prisma | 0.4 | 2700 | 3.35 | 882 | 9 | 320 x 260 x 208 |
| M22 | Trio | 0.4 | 2700 | 3.35 | 850 | 9 | 320 x 260 x 208 |

For each subject (n = 13), the scanner, voxel size (mm), TR (ms), TE (ms), TI (ms), flip angle (°), and dimensions of each T1w scan. For each subject, between 10 and 15 scans were taken and averaged to create a high-resolution image.

To correct for EPI distortions caused by magnetic field inhomogeneity, high-resolution field maps (0.6 × 0.6 × 0.7 mm voxel size), consisting of two GRE images with different TEs, were acquired from the 3T Siemens PRISMA scanner for M38 during a separate session under anaesthesia (TR = 917 ms, TE1 = 6.48 ms, TE2 = 8.94 ms, α = 55°, matrix size 140 × 140 × 66). The same field maps were also acquired for subjects 24 and 25, but in the same session as the experiments, while the monkey was performing a fixation task.

#### Experimental design and stimuli

Two phase-encoded retinotopic mapping experiments were performed to map different regions of the visual field^11,13,29^ Stimuli consisted of clockwise/counterclockwise rotating wedges and expanding/contracting rings. These stimuli covered a radius of 0.25°-12.5°. Wedge and ring stimuli consisted of apertures superimposed with expressive monkey faces^11^ or walking human figures^29^. The size of the wedges and rings increased with eccentricity to account for cortical magnification^35^. The moving ring and wedge apertures were used to elicit phase-locked activations, allowing for the calculation of eccentricity and polar angle representations, respectively. Opposing movement directions (expanding-contracting and clockwise-counterclockwise) were used to cancel phase errors caused by hemodynamic response lags.

The experimental design was adapted for the scanning parameters (TR) achieved for each subject. For the three subjects (M25, M38, and M24) in whom we acquired 0.6 mm isotropic voxels, the TR was 3000 ms. In this case, one full cycle of the wedge or ring stimuli consisted of 96 s. For all other subjects, the TR was 2000 ms, and each stimulus cycle lasted for 64 s. A Barco LCD projector was used to project the stimuli at 1400 x 1050 resolution and 60 Hz refresh rate onto a translucent screen located 57 cm from the subjects’ eyes.

Regardless of TR, each run comprised 4 cycles of either the ring or the wedge stimuli moving in one of the two directions, resulting in a total of 8 stimulus-sequence types. For these retinotopic mapping experiments, we used a purely exponential annuli-progression timing to compensate for the (exponential) magnification factor of the visual cortex.

The wedges were 45° wide, and the eccentricity annuli were scaled according to a log(r) law, to adapt the radial extent to the cortical magnification factor ^2^. In addition, aspect ratios of the figures superimposed onto the wedges and annuli were also held constant over the entire range of eccentricities by adjusting the size of the displayed animated object according to the same log(r) law. The breadths of the polar-angle wedges in the azimuthal direction and the lengths of the expanding annuli in the radial direction were specifically tailored to illuminate a given point on the screen for 8 s, the time required for the hemodynamic response to reach its maximum ^81^.

### 3. Quantification and Statistical Analysis

#### Data analysis

##### fMRI image reconstruction and preprocessing

The same preprocessing procedures as in Zhu et al., and Li et al.,^11,79^ were performed. To decrease artefacts caused by the monkey’s movements, the “optimized generalized autocalibration partially parallel acquisitions” (optimized GRAPPA) reconstruction method was used to reconstruct the EPI images. We used Bioimage suite and FreeSurfer for preprocessing, including skull stripping, slice-time correction, and motion correction with 6 degrees of freedom.

##### General linear model (GLM) analysis

Retinotopic maps (polar angle and eccentricity) were extracted following the procedure suggested by Sereno et al.,^2^ using Freesurfer. Nuisance regressors for the GLM analysis included three principal components derived from six motion-correction parameters and the linear and quadratic trend removals.

##### Surface segmentation, reconstruction, and projection

The same procedure as in Zhu et al.,^11^ was used to register each EPI template and the results in the same space as the reference anatomy of each subject. For the subjects acquired with 0.6 mm isotropic voxels, field maps were first used to correct EPI distortions caused by magnetic field inhomogeneity. GRE images acquired in each session were used as an intermediary to register between each functional session and the high-resolution template. The final results were then projected onto the surface by only taking voxels located between 30% and 70% of the cortical depth between the grey-white matter boundary and the pial surface. Values in these voxels were sampled in steps of 10% along the cortical thickness (depth sampling) and averaged to create the surface maps.

##### Field Sign Maps

Visual field sign maps were calculated in FreeSurfer in each subject’s individual surface space following the procedure described by Sereno et al.,^2,47^. Specifically, the clockwise angle (λ) between the eccentricity gradient (fovea to periphery) and polar angle gradient (LVF to HM to UVF) on the cortex was computed for each surface vertex. Vertices of the left hemisphere with an angle of approximately 90° (0 < λ < pi) were classified as a mirror image representation of the contralateral (right) hemifield, while vertices of the same hemisphere with an angle of approximately 270° (pi < λ < 2*pi) were classified as a non-mirror image representation of the same hemifield. For vertices of the right hemisphere, mirror and non-mirror image representations were determined vice versa.

##### Probabilistic Mapping

For each of the 13 subjects, the areas described above were delineated based on their retinotopic organization (V3A, V3d, and DLP were defined only in M25/38/24). These labels were then registered to a common template, in this case, the F99 macaque anatomical template ^82^, using surface-to-surface registration ^76^. By overlapping labels for each area per subject, a probabilistic representation of the area could be generated by following a percentage definition, i.e., each vertex was defined by the percentage of the 13 subjects that contained the relevant label. For V3A, V3d, and DLP, areas defined in one hemisphere were projected onto the other via surface-to-surface registration, and probabilistic maps were calculated from the pooled data of the six hemispheres. We used a 50% cut-off for the final area parcellation.

##### Transformation to MEBRAINS space

For each label/subject, data was transformed from the F99 template to the MEBRAINS template via surface registration. Iso-eccentricity and iso-polar lines were calculated in MEBRAINS surface space using the PyVista library for Python ^83^.

##### Overlap metrics

For each area, we calculated the total number of vertices included in the final parcellation (50% probability map). This number was then compared to the median number of vertices in agreement across subjects. The standard deviation was calculated across subjects.

## References

1 Vanduffel, W., and Zhu, Q. (2025). Comparative retinotopic mapping in macaques and humans. In Encyclopedia of the Human Brain (Elsevier), pp. 532–545. 10.1016/B978-0-12-820480-1.00199-6.

2. Sereno, M.I., Dale, A.M., Reppas, J.B., Kwong, K.K., Belliveau, J.W., Brady, T.J., Rosen, B.R., and Tootell, R.B.H. (1995). Borders of Multiple Visual Areas in Humans Revealed by Functional Magnetic Resonance Imaging. Science (1979). 268, 889–893. 10.1126/science.7754376.

3. Sereno, M.I., McDonald, C.T., and Allman, J.M. (1994). Analysis of Retinotopic Maps in Extrastriate Cortex. Cerebral Cortex 4, 601–620. 10.1093/cercor/4.6.601.

4. Wade, A.R., Brewer, A.A., Rieger, J.W., and Wandell, B.A. (2002). Functional measurements of human ventral occipital cortex: retinotopy and colour. Philos. Trans. R. Soc. Lond. B Biol. Sci. 357, 963–973. 10.1098/rstb.2002.1108.

5. Tootell, R.B.H., Hadjikhani, N.K., Vanduffel, W., Liu, A.K., Mendola, J.D., Sereno, M.I., and Dale, A.M. (1998). Functional analysis of primary visual cortex (V1) in humans. Proceedings of the National Academy of Sciences 95, 811–817. 10.1073/pnas.95.3.811.

6. Wang, L., Mruczek, R.E.B., Arcaro, M.J., and Kastner, S. (2015). Probabilistic Maps of Visual Topography in Human Cortex. Cerebral Cortex 25, 3911–3931. 10.1093/cercor/bhu277.

7. Gattass, R., Sousa, A.P.B., and Gross, C.G. (1988). Visuotopic organization and extent of V3 and V4 of the macaque. Journal of Neuroscience 8, 1831–1845. 10.1523/jneurosci.08-06-01831.1988.

8. Felleman, D.J., and Van Essen, D.C. (1991). Distributed Hierarchical Processing in the Primate Cerebral Cortex. Cerebral Cortex 1, 1–47. 10.1093/cercor/1.1.1-a.

9. Arcaro, M.J., and Livingstone, M.S. (2017). Retinotopic Organization of Scene Areas in Macaque Inferior Temporal Cortex. The Journal of Neuroscience 37, 7373–7389. 10.1523/JNEUROSCI.0569-17.2017.

10. Rima, S., Cottereau, B.R., Héjja-Brichard, Y., Trotter, Y., and Durand, J.-B. (2020). Wide-field retinotopy reveals a new visuotopic cluster in macaque posterior parietal cortex. Brain Struct. Funct. 225, 2447–2461. 10.1007/s00429-020-02134-2.

11. Zhu, Q., and Vanduffel, W. (2019). Submillimeter fMRI reveals a layout of dorsal visual cortex in macaques, remarkably similar to New World monkeys. Proceedings of the National Academy of Sciences 116, 2306–2311. 10.1073/pnas.1805561116.

12. Sepe, A., Panormita, M., Zhu, Q., Li, X., Leopold, D.A., Tamietto, M., Bonini, L., and Vanduffel, W. (2025). Lateralized visuotopic organization in the macaque superior colliculus revealed by fMRI. Prog. Neurobiol. 254, 102842. 10.1016/j.pneurobio.2025.102842.

13. Kolster, H., Janssens, T., Orban, G.A., and Vanduffel, W. (2014). The Retinotopic Organization of Macaque Occipitotemporal Cortex Anterior to V4 and Caudoventral to the Middle Temporal (MT) Cluster. Journal of Neuroscience 34, 10168–10191. 10.1523/JNEUROSCI.3288-13.2014.

14. Abdollahi, R.O., Kolster, H., Glasser, M.F., Robinson, E.C., Coalson, T.S., Dierker, D., Jenkinson, M., Van Essen, D.C., and Orban, G.A. (2014). Correspondences between retinotopic areas and myelin maps in human visual cortex. Neuroimage 99, 509–524. 10.1016/j.neuroimage.2014.06.042.

15. Rosa, M.G.. P., Piñon, M.C., Gattass, R., and Sousa, A.P.B. (2000). Third tier ventral extrastriate cortex in the New World monkey, Cebus apella. Exp. Brain Res. 132, 287–305. 10.1007/s002210000344.

16. Kaas, J.H., Roe, A.W., Baldwin, M.K.L., and Lyon, D.C. (2015). Resolving the organization of the territory of the third visual area: A new proposal. Vis. Neurosci. 32, E016. 10.1017/S0952523815000152.

17. Messinger, A., Jung, B., Sponheim, C., and Ungerleider, L.G. (2019). fMRI mapping of retinotopy using face and object stimuli in rhesus monkeys. J. Vis. 19, 261a. 10.1167/19.10.261a.

18. DeSimone, K., Viviano, J.D., and Schneider, K.A. (2015). Population Receptive Field Estimation Reveals New Retinotopic Maps in Human Subcortex. Journal of Neuroscience 35, 9836–9847. 10.1523/JNEUROSCI.3840-14.2015.

19. Qian, M., Wang, J., Gao, Y., Chen, M., Liu, Y., Zhou, D., Lu, H.D., Zhang, X., Hu, J.M., and Roe, A.W. (2025). Multiple loci for foveolar vision in macaque monkey visual cortex. Nat. Neurosci. 28, 137–149. 10.1038/s41593-024-01810-4.

20. Fize, D., Vanduffel, W., Nelissen, K., Denys, K., d’Hotel, C.C., Faugeras, O., and Orban, G.A. (2003). The Retinotopic Organization of Primate Dorsal V4 and Surrounding Areas: A Functional Magnetic Resonance Imaging Study in Awake Monkeys. The Journal of Neuroscience 23, 7395–7406. 10.1523/JNEUROSCI.23-19-07395.2003.

21. Paxinos, G., Huang, X.-F., Petrides, M., and Toga, A.W. (2009). The Rhesus Monkey Brain in Stereotaxic Coordinates 2nd ed. (Elsevier Science).

22. Saleem, K.S.., and Logothetis, Nikos. (2012). A combined MRI and histology atlas of the rhesus monkey brain in stereotaxic coordinates (Elsevier/AP).

23. Amunts, K., and Zilles, K. (2001). ADVANCES IN CYTOARCHITECTONIC MAPPING OF THE HUMAN CEREBRAL CORTEX. Neuroimaging Clin. N. Am. 11, 151–169. 10.1016/S1052-5149(25)00698-7.

24. Amunts, K., Mohlberg, H., Bludau, S., and Zilles, K. (2020). Julich-Brain: A 3D probabilistic atlas of the human brain’s cytoarchitecture. Science (1979). 369, 988–992. 10.1126/science.abb4588.

25. Sarubbo, S., Tate, M., De Benedictis, A., Merler, S., Moritz-Gasser, S., Herbet, G., and Duffau, H. (2020). Mapping critical cortical hubs and white matter pathways by direct electrical stimulation: an original functional atlas of the human brain. Neuroimage 205, 116237. 10.1016/j.neuroimage.2019.116237.

26. Liu, F., Zhang, Z., Chen, Y., Wei, L., Xu, Y., Li, Z., and Zhu, C. (2023). MNI2CPC: A probabilistic cortex-to-scalp mapping for non-invasive brain stimulation targeting. Brain Stimul. 16, 1733–1742. 10.1016/j.brs.2023.11.011.

27. Olchanyi, M.D., Schreier, D.R., Li, J., Maffei, C., Sorby-Adams, A., Kinney, H.C., Healy, B.C., Freeman, H.J., Shless, J., Destrieux, C., et al. (2026). Probabilistic mapping and automated segmentation of human brainstem white matter bundles. Proceedings of the National Academy of Sciences 123. 10.1073/pnas.2509321123.

28. Van Essen, D.C., and Dierker, D.L. (2007). Surface-Based and Probabilistic Atlases of Primate Cerebral Cortex. Neuron 56, 209–225. 10.1016/j.neuron.2007.10.015.

29. Janssens, T., Zhu, Q., Popivanov, I.D., and Vanduffel, W. (2014). Probabilistic and Single-Subject Retinotopic Maps Reveal the Topographic Organization of Face Patches in the Macaque Cortex. Journal of Neuroscience 34, 10156–10167. 10.1523/JNEUROSCI.2914-13.2013.

30. Dworetsky, A., Seitzman, B.A., Adeyemo, B., Neta, M., Coalson, R.S., Petersen, S.E., and Gratton, C. (2021). Probabilistic mapping of human functional brain networks identifies regions of high group consensus. Neuroimage 237, 118164. 10.1016/j.neuroimage.2021.118164.

31. Benson, N.C., and Winawer, J. (2018). Bayesian analysis of retinotopic maps. Elife 7. 10.7554/eLife.40224.

32. Balan, P.F., Zhu, Q., Li, X., Niu, M., Rapan, L., Funck, T., Wang, H., Bakker, R., Palomero-Gallagher, N., and Vanduffel, W. (2024). MEBRAINS 1.0: A new population-based macaque atlas. Imaging Neuroscience 2. 10.1162/imag_a_00077.

33. Fischl, B., Sereno, M.I., Tootell, R.B.H., and Dale, A.M. (1999). High-resolution intersubject averaging and a coordinate system for the cortical surface. Hum. Brain Mapp. 8, 272–284. 10.1002/(SICI)1097-0193(1999)8:4<272::AID-HBM10>3.0.CO;2-4.

34. Buckner, R.L., and Yeo, B.T.T. (2014). Borders, map clusters, and supra-areal organization in visual cortex. Neuroimage 93, 292–297. 10.1016/j.neuroimage.2013.12.036.

35. Daniel, P.M., and Whitteridge, D. (1961). The representation of the visual field on the cerebral cortex in monkeys. J. Physiol. 159, 203–221. 10.1113/jphysiol.1961.sp006803.

36. Gattass, R., Gross, C.G., and Sandell, J.H. (1981). Visual topography of V2 in the macaque. Journal of Comparative Neurology 201, 519–539. 10.1002/cne.902010405.

37. Wandell, B.A., Dumoulin, S.O., and Brewer, A.A. (2007). Visual Field Maps in Human Cortex. Neuron 56, 366–383. 10.1016/j.neuron.2007.10.012.

38. Benson, N.C., Butt, O.H., Brainard, D.H., and Aguirre, G.K. (2014). Correction of Distortion in Flattened Representations of the Cortical Surface Allows Prediction of V1-V3 Functional Organization from Anatomy. PLoS Comput. Biol. 10, e1003538. 10.1371/journal.pcbi.1003538.

39. Brewer, A.A., Press, W.A., Logothetis, N.K., and Wandell, B.A. (2002). Visual Areas in Macaque Cortex Measured Using Functional Magnetic Resonance Imaging. The Journal of Neuroscience 22, 10416–10426. 10.1523/JNEUROSCI.22-23-10416.2002.

40. Markov, N.T., Misery, P., Falchier, A., Lamy, C., Vezoli, J., Quilodran, R., Gariel, M.A., Giroud, P., Ercsey-Ravasz, M., Pilaz, L.J., et al. (2011). Weight Consistency Specifies Regularities of Macaque Cortical Networks. Cerebral Cortex 21, 1254–1272. 10.1093/cercor/bhq201.

41. Reveley, C., Gruslys, A., Ye, F.Q., Glen, D., Samaha, J., E. Russ, B., Saad, Z., K. Seth, A., Leopold, D.A., and Saleem, K.S. (2016). Three-Dimensional Digital Template Atlas of the Macaque Brain. Cerebral Cortex. 10.1093/cercor/bhw248.

42. Lewis, J.W., and Van Essen, D.C. (2000). Mapping of architectonic subdivisions in the macaque monkey, with emphasis on parieto-occipital cortex. J. Comp. Neurol. 428, 79–111. 10.1002/1096-9861(20001204)428:1<79::AID-CNE7>3.0.CO;2-Q.

43. Van Essen, D.C., Newsome, W.T., and Maunsell, J.H.R. (1984). The visual field representation in striate cortex of the macaque monkey: Asymmetries, anisotropies, and individual variability. Vision Res. 24, 429–448. 10.1016/0042-6989(84)90041-5.

44. Felleman, D.J., and Van Essen, D.C. (1987). Receptive field properties of neurons in area V3 of macaque monkey extrastriate cortex. J. Neurophysiol. 57, 889–920. 10.1152/jn.1987.57.4.889.

45. Tootell, R.B.H., Mendola, J.D., Hadjikhani, N.K., Ledden, P.J., Liu, Arthur. K., Reppas, J.B., Sereno, M.I., and Dale, A.M. (1997). Functional Analysis of V3A and Related Areas in Human Visual Cortex. The Journal of Neuroscience 17, 7060–7078. 10.1523/JNEUROSCI.17-18-07060.1997.

46. Essen, D.C., and Zeki, S.M. (1978). The topographic organization of rhesus monkey prestriate cortex. J. Physiol. 277, 193–226. 10.1113/jphysiol.1978.sp012269.

47. Sereno, M.I., McDonald, C.T., and Allman, J.M. (2015). Retinotopic organization of extrastriate cortex in the owl monkey—dorsal and lateral areas. Vis. Neurosci. 32, E021. 10.1017/S0952523815000206.

48. Gattass, R., Lima, B., Soares, J.G.M., and Ungerleider, L.G. (2015). Controversies about the visual areas located at the anterior border of area V2 in primates. Vis. Neurosci. 32, E019. 10.1017/S0952523815000188.

49. Zeki, S.M. (1973). Colour coding in rhesus monkey prestriate cortex. Brain Res. 53, 422–427. 10.1016/0006-8993(73)90227-8.

50. Shipp, S., and Zeki, S. (1985). Segregation of pathways leading from area V2 to areas V4 and V5 of macaque monkey visual cortex. Nature 315, 322–324. 10.1038/315322a0.

51. Roe, A.W., Chelazzi, L., Connor, C.E., Conway, B.R., Fujita, I., Gallant, J.L., Lu, H., and Vanduffel, W. (2012). Toward a Unified Theory of Visual Area V4. Neuron 74, 12–29. 10.1016/j.neuron.2012.03.011.

52. Maguire, W., and Baizer, J. (1984). Visuotopic organization of the prelunate gyrus in rhesus monkey. The Journal of Neuroscience 4, 1690–1704. 10.1523/JNEUROSCI.04-07-01690.1984.

53. Zeki, S.M. (1977). Colour coding in the superior temporal sulcus of rhesus monkey visual cortex. Proc. R. Soc. Lond. B Biol. Sci. 197, 195–223. 10.1098/rspb.1977.0065.

54. Zeki, S.M. (1971). Cortical projections from two prestriate areas in the monkey. Brain Res. 34, 19–35. 10.1016/0006-8993(71)90348-9.

55. Pigarev, I.N., Nothdurft, H.-C., and Kastner, S. (2002). Neurons with radial receptive fields in monkey area V4A: evidence of a subdivision of prelunate gyrus based on neuronal response properties. Exp. Brain Res. 145, 199–206. 10.1007/s00221-002-1112-y.

56. Stepniewska, I., Collins, C.E., and Kaas, J.H. (2005). Reappraisal of DL/V4 Boundaries Based on Connectivity Patterns of Dorsolateral Visual Cortex in Macaques. Cerebral Cortex 15, 809–822. 10.1093/cercor/bhh182.

57. Kolster, H., Mandeville, J.B., Arsenault, J.T., Ekstrom, L.B., Wald, L.L., and Vanduffel, W. (2009). Visual Field Map Clusters in Macaque Extrastriate Visual Cortex. The Journal of Neuroscience 29, 7031–7039. 10.1523/JNEUROSCI.0518-09.2009.

58. Kolster, H., Peeters, R., and Orban, G.A. (2010). The Retinotopic Organization of the Human Middle Temporal Area MT/V5 and Its Cortical Neighbors. Journal of Neuroscience 30, 9801–9820. 10.1523/JNEUROSCI.2069-10.2010.

59. Zeki, S.M. (1974). Functional organization of a visual area in the posterior bank of the superior temporal sulcus of the rhesus monkey. J. Physiol. 236, 549–573. 10.1113/jphysiol.1974.sp010452.

60. Gattass, R., and Gross, C.G. (1981). Visual topography of striate projection zone (MT) in posterior superior temporal sulcus of the macaque. J. Neurophysiol. 46, 621–638. 10.1152/jn.1981.46.3.621.

61. Tanaka, K., and Saito, H. (1989). Analysis of motion of the visual field by direction, expansion/contraction, and rotation cells clustered in the dorsal part of the medial superior temporal area of the macaque monkey. J. Neurophysiol. 62, 626–641. 10.1152/jn.1989.62.3.626.

62. Komatsu, H., and Wurtz, R.H. (1988). Relation of cortical areas MT and MST to pursuit eye movements. I. Localization and visual properties of neurons. J. Neurophysiol. 60, 580–603. 10.1152/jn.1988.60.2.580.

63. Maunsell, J., and van Essen, D. (1983). The connections of the middle temporal visual area (MT) and their relationship to a cortical hierarchy in the macaque monkey. The Journal of Neuroscience 3, 2563–2586. 10.1523/JNEUROSCI.03-12-02563.1983.

64. Desimone, R., and Ungerleider, L.G. (1986). Multiple visual areas in the caudal superior temporal sulcus of the macaque. Journal of Comparative Neurology 248, 164–189. 10.1002/cne.902480203.

65. Boussaoud, D., Desimone, R., and Ungerleider, L.G. (1991). Visual topography of area TEO in the macaque. Journal of Comparative Neurology 306, 554–575. 10.1002/cne.903060403.

66. Ungerleider, L.G., Galkin, T.W., Desimone, R., and Gattass, R. (2008). Cortical Connections of Area V4 in the Macaque. Cerebral Cortex 18, 477–499. 10.1093/cercor/bhm061.

67. Arcaro, M.J., Pinsk, M.A., Li, X., and Kastner, S. (2011). Visuotopic Organization of Macaque Posterior Parietal Cortex: A Functional Magnetic Resonance Imaging Study. The Journal of Neuroscience 31, 2064–2078. 10.1523/JNEUROSCI.3334-10.2011.

68. Ben Hamed, S., Duhamel, J.-R., Bremmer, F., and Graf, W. (2001). Representation of the visual field in the lateral intraparietal area of macaque monkeys: a quantitative receptive field analysis. Exp. Brain Res. 140, 127–144. 10.1007/s002210100785.

69. Blatt, G.J., Andersen, R.A., and Stoner, G.R. (1990). Visual receptive field organization and cortico-cortical connections of the lateral intraparietal area (area LIP) in the macaque. Journal of Comparative Neurology 299, 421–445. 10.1002/cne.902990404.

70. Patel, G.H., Shulman, G.L., Baker, J.T., Akbudak, E., Snyder, A.Z., Snyder, L.H., and Corbetta, M. (2010). Topographic organization of macaque area LIP. Proceedings of the National Academy of Sciences 107, 4728–4733. 10.1073/pnas.0908092107.

71. Jeffs, J., Federer, F., and Angelucci, A. (2015). Corticocortical connection patterns reveal two distinct visual cortical areas bordering dorsal V2 in marmoset monkey. Vis. Neurosci. 32, E012. 10.1017/S0952523815000097.

72. Li, X., Zhu, Q., and Vanduffel, W. (2021). Myelin densities in retinotopically defined dorsal visual areas of the macaque. Brain Struct. Funct. 226, 2869–2880. 10.1007/s00429-021-02363-z.

73. Kastner, S., Chen, Q., Jeong, S.K., and Mruczek, R.E.B. (2017). A brief comparative review of primate posterior parietal cortex: A novel hypothesis on the human toolmaker. Neuropsychologia 105, 123–134. 10.1016/j.neuropsychologia.2017.01.034.

74. Jung, B., Taylor, P.A., Seidlitz, J., Sponheim, C., Perkins, P., Ungerleider, L.G., Glen, D., and Messinger, A. (2021). A comprehensive macaque fMRI pipeline and hierarchical atlas. Neuroimage 235, 117997. 10.1016/j.neuroimage.2021.117997.

75. Boussaoud, D., Ungerleider, L.G., and Desimone, R. (1990). Pathways for motion analysis: Cortical connections of the medial superior temporal and fundus of the superior temporal visual areas in the macaque. Journal of Comparative Neurology 296, 462–495. 10.1002/cne.902960311.

76. Pantazis, D., Joshi, A., Jiang, J., Shattuck, D.W., Bernstein, L.E., Damasio, H., and Leahy, R.M. (2010). Comparison of landmark-based and automatic methods for cortical surface registration. Neuroimage 49, 2479–2493. 10.1016/j.neuroimage.2009.09.027.

77. Froudist-Walsh, S., Niu, M., Rapan, L., and Palomero-Gallagher, N. (2023). Neurotransmitter receptor densities per neuron across macaque cortex (v1.0). Preprint at EBRAINS, 10.25493/5HK3-S8M 10.25493/5HK3-S8M.

78. Vanduffel, W., Fize, D., Mandeville, J.B., Nelissen, K., Van Hecke, P., Rosen, B.R., Tootell, R.B.H., and Orban, G.A. (2001). Visual Motion Processing Investigated Using Contrast Agent-Enhanced fMRI in Awake Behaving Monkeys. Neuron 32, 565–577. 10.1016/S0896-6273(01)00502-5.

79. Li, X., Zhu, Q., Janssens, T., Arsenault, J.T., and Vanduffel, W. (2019). In Vivo Identification of Thick, Thin, and Pale Stripes of Macaque Area V2 Using Submillimeter Resolution (f)MRI at 3 T. Cerebral Cortex 29, 544–560. 10.1093/cercor/bhx337.

80. Janssens, T., Keil, B., Farivar, R., McNab, J.A., Polimeni, J.R., Gerits, A., Arsenault, J.T., Wald, L.L., and Vanduffel, W. (2012). An implanted 8-channel array coil for high-resolution macaque MRI at 3T. Neuroimage 62, 1529–1536. 10.1016/j.neuroimage.2012.05.028.

81. Leite, F.P., and Mandeville, J.B. (2006). Characterization of event-related designs using BOLD and IRON fMRI. Neuroimage 29, 901–909. 10.1016/j.neuroimage.2005.08.022.

82. Van Essen, D.C. (2002). Surface-Based Atlases of Cerebellar Cortex in the Human, Macaque, and Mouse. Ann. N. Y. Acad. Sci. 978, 468–479. 10.1111/j.1749-6632.2002.tb07588.x.

83. Sullivan, C., and Kaszynski, A. (2019). PyVista: 3D plotting and mesh analysis through a streamlined interface for the Visualization Toolkit (VTK). J. Open Source Softw. 4, 1450. 10.21105/joss.01450.

