## Supplemental Document for "Probabilistic Retinotopic Parcellation of the Macaque Visual Cortex"

| ROI | LH median (%) | LH std | RH median (%) | RH std |
|---|---|---|---|---|
| CIP1 | 43.21 | 19.61 | 45.83 | 23.88 |
| CIP2 | 45.93 | 17.62 | 48.82 | 22.89 |
| FST | 53.6 | 14.32 | 55.71 | 13.02 |
| LIPvt | 68.8 | 17.07 | 70.85 | 13.36 |
| MST | 44.04 | 22.74 | 39.92 | 21.07 |
| MT | 65.05 | 15.37 | 70 | 14.31 |
| OTd | 48.26 | 20.72 | 53.79 | 11.67 |
| PITd | 55.36 | 20.09 | 66.18 | 13.06 |
| PITv | 66.54 | 16.64 | 73.73 | 10.94 |
| V1 | 91.75 | 3.82 | 91.64 | 6.82 |
| V2d | 84.24 | 6.39 | 82.15 | 7.07 |
| V2v | 89.17 | 4.26 | 86.76 | 4.87 |
| V3v | 71.51 | 10.77 | 72.33 | 11.31 |
| V4 | 71.23 | 11.29 | 76.19 | 8.92 |
| V4t | 50.88 | 17.2 | 46.45 | 16.79 |
| V4A | 47.67 | 14.53 | 55.7 | 11.89 |

**Table S1:** Ratio of overlapped vertices between subjects and final parcellation. For each area, we calculated the total number of vertices included in the final parcellation (50% probability map). This number was then compared to the median number of vertices in agreement across subjects. Standard deviation was calculated across subjects. V3A, V3d, and DLP are excluded from this calculation as for these areas definitions in one hemisphere were projected onto the other via surface-to-surface registration, and probabilistic maps were calculated from the pooled data of the six hemispheres.

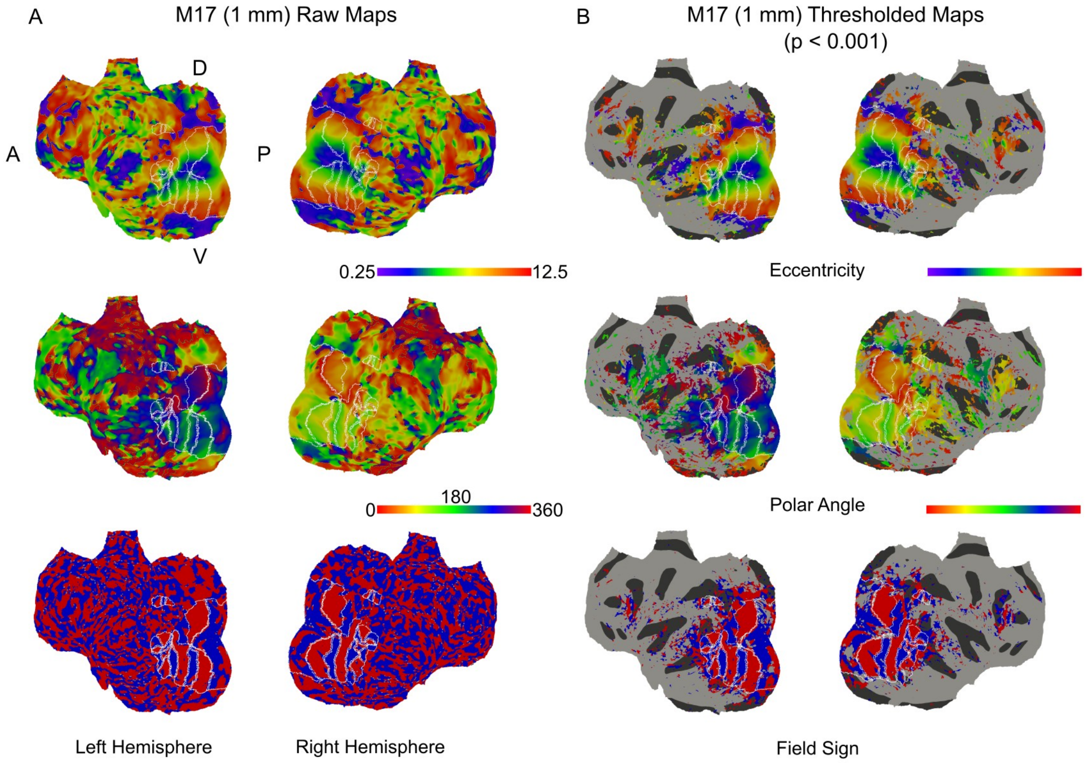


**Figure S1:** Example raw data from a single subject (M17). Each row shows flatmap representations of the MEBRAINS template with the eccentricity, polar angle, and field sign maps for one subject. The left set of maps is unthresholded, and the right set of maps is thresholded to a p-value < 0.001. Color bars below each set of maps.

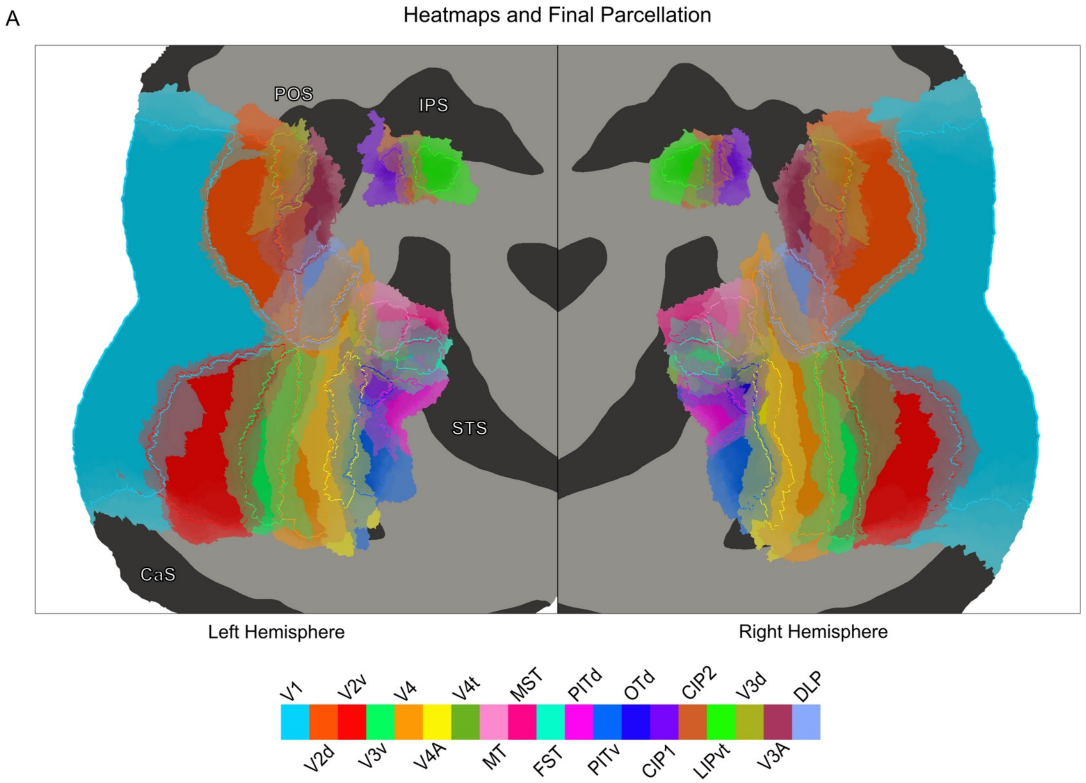


**Figure S2:** (A) Probabilistic heatmaps for each area in our final parcellation overlaid on the left and right flatmap representations of the MEBRAINS template. Sulcus names are shown in white with black-outlined text. Colormap show at the bottom of the figure.
